# Spontaneous human CD8 T cell and EAE-inducible human CD4/CD8 T cell lesions in the brain and spinal cord of HLA-DRB1*15-positive multiple sclerosis PBMC humanized mice

**DOI:** 10.1101/2023.06.07.543414

**Authors:** Irini Papazian, Maria Kourouvani, Anastasia Dagkonaki, Vasileios Gouzouasis, Lila Dimitrakopoulou, Nikolaos Markoglou, Fotis Badounas, Theodore Tselios, Maria Anagnostouli, Lesley Probert

**Author notes:** Correspondence: Lesley Probert.

## Abstract

Autoimmune diseases of the central nervous system (CNS) such as multiple sclerosis (MS) are only partially represented in current experimental models and the development of humanized immune mice is crucial for better understanding of immunopathogenesis and testing of therapeutics. We describe a humanized mouse model with several key features of MS. Severely immunodeficient B2m-NOG mice were transplanted with peripheral blood mononuclear cells (PBMC) from HLA-DRB1-typed MS and healthy (HI) donors and showed rapid engraftment by human T and B lymphocytes. Mice receiving cells from MS patients with recent/ongoing Epstein-Barr virus (EBV) reactivation showed high B cell engraftment capacity. Both HLA-DRB1*15 (DR15) MS and DR15 HI mice, not HLA-DRB1*13 MS mice, developed human T cell infiltration of CNS borders and parenchyma. DR15 MS mice uniquely developed inflammatory lesions in brain and spinal cord grey matter, with spontaneous, hCD8 T cell lesions, and mixed hCD8/hCD4 T cell lesions in EAE immunized mice, with variation in localization and severity between different patient donors. Main limitations of this model for further development are poor monocyte engraftment and lack of demyelination, lymph node organization and IgG responses. These results show that PBMC humanized mice represent promising research tools for investigating MS immunopathology in a patient-specific approach.

## Introduction

Multiple sclerosis (MS) is a complex chronic inflammatory demyelinating and neurodegenerative disease of the central nervous system (CNS) with autoimmune characteristics. Pathological hallmarks are focal plaques of immune-mediated demyelination in the white matter, inflammation at CNS borders and diffuse demyelination and neurodegeneration of the grey and white matter of the brain and spinal cord. Genome wide association studies show that the vast majority of MS risk genes are related to immune activation and function, thereby strongly supporting the role of adaptive immunity in MS pathogenesis. These include the strongest MS associated genetic risk factor, human leukocyte antigen (HLA) DRB1*15 haplotype, encoded by two DRB* genes, *DRB1*\**15:01* (DR2b) and *DRB5*01:01* (DR2a), of the MHC class II (MHCII) that restricts CD4^+^ T cells (1,2). Pathological analyses of autopsy tissues reveal CD8^+^ lymphocytes and monocytes as the main CNS infiltrating cells, with B cells accumulating in meningeal and perivascular spaces, and relatively few infiltrating CD4^+^ T cells (3). Studies in animal models, mainly experimental autoimmune encephalomyelitis (EAE), showing that myelin-reactive T cells together with peripheral monocytes are necessary for EAE induction, as well as the effectiveness of current immunotherapies in MS patients that work mainly by targeting peripheral immune responses, further support an autoimmune pathogenesis for MS.

EAE and MS share many clinical, immunological, and pathological similarities (4,5), but they also present significant differences (6,7). One difference is the predominance of CD8^+^ T cells in MS lesions, while these cells are scarce in the CNS of EAE mice (8). In addition, the inflammatory process in EAE primarily targets the spinal cord, whereas in MS it also targets the brain. Moreover, while some MS drugs were discovered from research in the EAE model (natalizumab, glatiramer acetate), there is a relatively low or moderate success rate of drugs that showed high efficacy in both the EAE model and the human disease (9). These differences infer that different pathways or mechanisms are prevalent in MS versus EAE (10,11). Differences are further highlighted by recent clinical data showing the critical importance of B cells to MS pathology, a cell population that does not show important functional participation in most EAE models (12,13). The development of animal models capable of developing a human immune system is crucial to the study of human immune-mediated diseases such as MS, and for evaluating the safety and efficacy of human immunotherapies. Most current studies with humanized mouse models use immunodeficient non-obese diabetic (NOD) - severe combined immunodeficiency disorder (*scid*) mice deficient or mutant for the common cytokine receptor γ-chain gene (*IL2rg)* (named NSG or NOG mice, respectively) that lack murine B, T cells and NK cells and have defective innate immune responses (14–17). These mouse strains readily engraft with human peripheral blood mononuclear cells (PBMC) without prior irradiation, as murine MHC molecules support the proliferation of both human CD4^+^ and CD8^+^ T cells, with CD8^+^ T cells predominating. Recent variants of these strains lacking MHC class I molecules (B2m) show increased CD4 to CD8 ratios and a longer time window before onset of xenogeneic graft versus host disease (GVHD) and represent improved strains for the study of human immune responses (18,19). Previous studies showed that PBMC humanized NSG mice with cells from healthy donors are susceptible to EAE, exhibiting subclinical disease, with infiltration of the brain and spinal cord by CD4^+^ and CD8^+^ T cells, but not monocytes (20).

To investigate the potential of PBMC from MS patients to induce CNS-directed immunopathology in humanized mice, we engrafted B2m-NOD/*scid*-IL2Rγ^null^ (B2m-NOG) mice with PBMC from human donors based on HLA-DRB1 genotype, RRMS diagnosis and therapy, and monitored CNS inflammation that developed spontaneously or after immunization for EAE with myelin antigens. We show that PBMC from MS and healthy (HI) donors rapidly engraft B2m-NOG mice with human (h) CD4^+^ and hCD8^+^ T cells and B cells, with donor-specific differences towards proportions of human immune cell populations engrafted and propensity to develop CNS inflammation. Mice transplanted with PBMC from HLA-DRB1*15-positive (DR15) MS patients uniquely developed spontaneous hCD8^+^ T cell lesions in the spinal cord grey matter and brainstem, and prominent brain white and grey matter lesions with mixed hCD4/hCD8 T cells in mice immunized with myelin peptides. Our results show that PBMC humanized B2m-NOG mice partially reproduce MS immunopathology and can be used to investigate human immune responses towards CNS in a simple, rapid, and personalized manner.

## Results

### Reconstitution of human adaptive immune system in B2m-NOG mice engrafted with PBMC from MS patient and HI donors

Six long-term RRMS patients, of which five were HLA-DRB1*15-positive (DR15 MS1-5) and one HLA-DRB1*13-positive (DR13 MS), all presenting with highly active disease following immunomodulatory treatment with natalizumab, were selected as PBMC donors (Table 1). A DR15-positive healthy individual (DR15 HI) was selected as a control donor. Fresh blood samples were used for immunoprofiling by flow cytometry with a panel of standard human immune cell marker antibodies (Supplementary Table 2A, B), for screening of plasma antibody responses to viruses (Table 1; Supplementary Figure 2C for clinical interpretation of Epstein-Barr virus/EBV results), and for the isolation of fresh PBMC for transplantation. Groups of 7-8 B2m-NOG mice were injected intravenously with freshly prepared PBMC (1 x 10^7^/mouse) from each donor (Supplementary Table 3, protocol 1). The transplanted mice were monitored for human immune cell engraftment by flow cytometry analysis of small blood samples recovered from the tail vein at different time points and all showed progressive engraftment by human CD45^+^ (hCD45^+^) leukocytes from day 7 (Fig. 1A; Supplementary Figure 1A). At day 14 post-transplantation (dpt 14), 4-5 mice from each group were immunized for EAE using a mixture of immunodominant T cell myelin peptide antigens, following two different protocols. In EAE experiment 1, groups of DR13 MS, DR15 MS1 and DR15 HI mice were immunized twice, 7 days apart, using 200 μg each peptide/mouse/injection. In EAE experiment 2, groups of DR15 MS2-5 were immunized once, using 100 μg each peptide/mouse. Several PBMC donors used in this study were previously shown to have human T cell proliferation responses to myelin peptides (21) (Table 1). Non-immunized mice, and mice immunized for EAE with low-dose peptides (1x100 μg) showed steadily increasing levels of blood hCD45^+^ leukocytes up to sacrifice at dpt 42 (Fig. 1A; Supplementary Figure 1A). Mice immunized for EAE with high-dose peptides (2x200 μg) showed reduced levels of blood hCD45^+^ leukocytes following immunization (Fig. 1A). Further FACS analysis showed preferential expansion of hCD4^+^ T lymphocytes compared to hCD8^+^ T lymphocytes in blood and spleens of all engrafted mice (Fig. 1B, C; Supplementary Figure 1B), a finding consistent with a previous report describing the B2m-NSG model (18).

**Figure 1.**
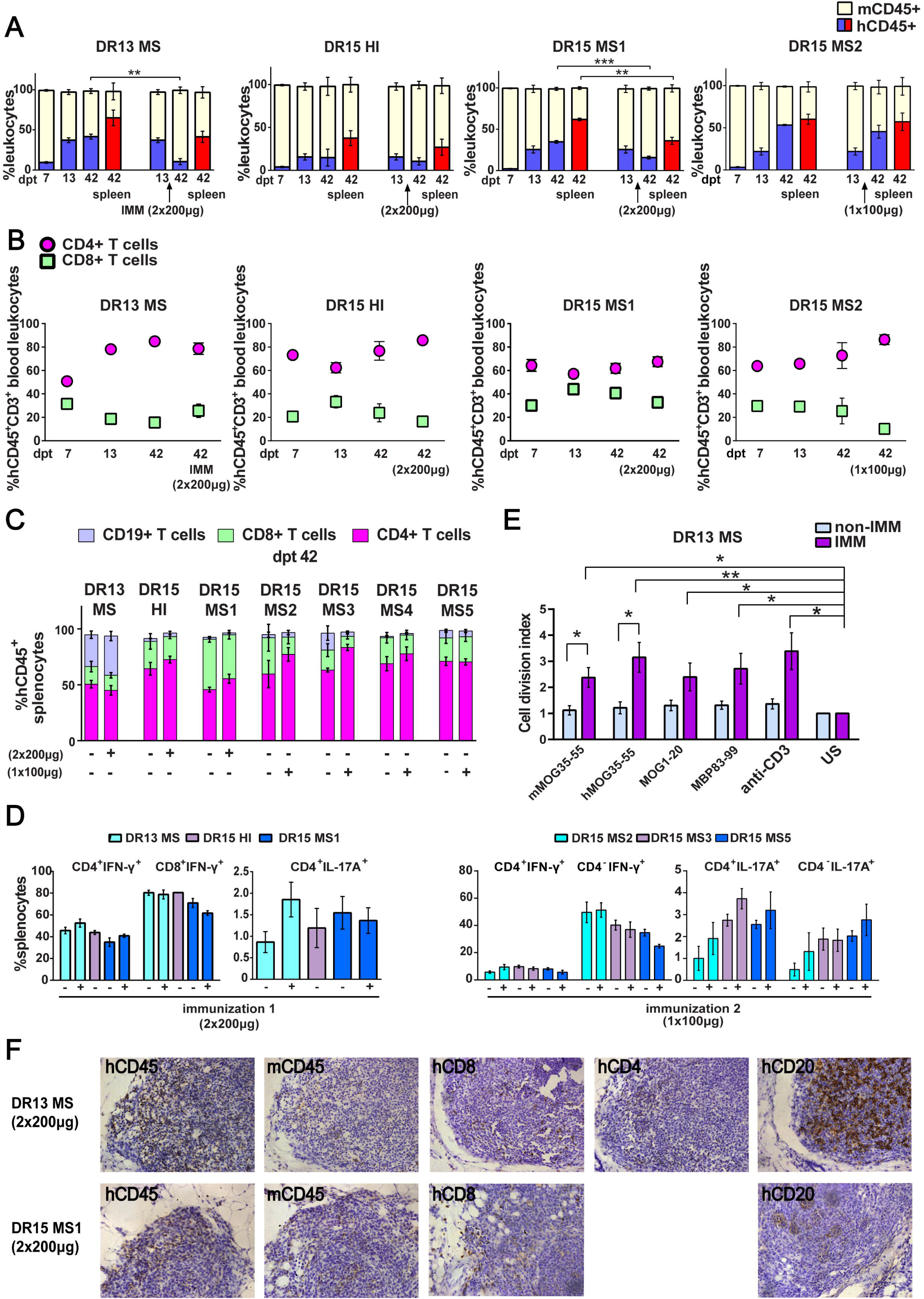
Reconstitution of a human adaptive immune system in B2m-NOG mice engrafted with PBMC from MS patient and healthy donors. **(A)** Progressive engraftment of human (h) CD45+ leukocytes from donors in groups of B2m-NOG mice, non-immunized (left) and immunized for EAE using a myelin peptide cocktail in repeat immunizations with 200 μg/each myelin peptide (EAE experiment 1), or single immunization with 100 μg/each myelin peptide (EAE experiment 2), measured in peripheral blood samples taken at different time points, and in spleen recovered at sacrifice 42 days’ post-transplantation (dpt 42). **(B)** Proportions of hCD4^+^ and hCD8^+^ T cells in blood hCD45^+^CD3^+^ T cells at different time points. **(C)** Proportions of human immune cell subpopulations in spleens of immunized and non-immunized mice at dpt 42. **(D)** Proportions of interferon-γ-producing and IL-17A-producing CD4^+^ and CD8^+^ (or CD4^-^) T cells in splenocytes recovered from mice in the different groups at dpt 42. **(E)** Antigen-specific T cell proliferation responses to the immunizing antigens (mMOG35-55, hMOG35-55, MOG1-20, MBP83-99), anti-hCD3 (positive control) and medium (unstimulated; US), in splenocytes recovered from non-immunized and immunized DR13 MS PBMC humanized mice at dpt 42. Results are expressed as a cell division index. **(F)** Draining lymphoid structures in inguinal fat of mice engrafted with DR13 MS and DR15 MS PBMC and immunized with myelin peptides, recovered at dpt 42. Immunohistochemistry revealed the presence of hCD45- and mCD45-positive leukocytes, hCD4- and hCD8-positive T cells, and hCD20-positive B cells (latter only DR13 MS mice). All results are depicted as mean ± SEM. Statistical significance is shown after pairwise comparisons between groups using Student’s t test (*p ≤ 0.05, **p ≤ 0.01, ***p ≤ 0.001).

**Table 1.**
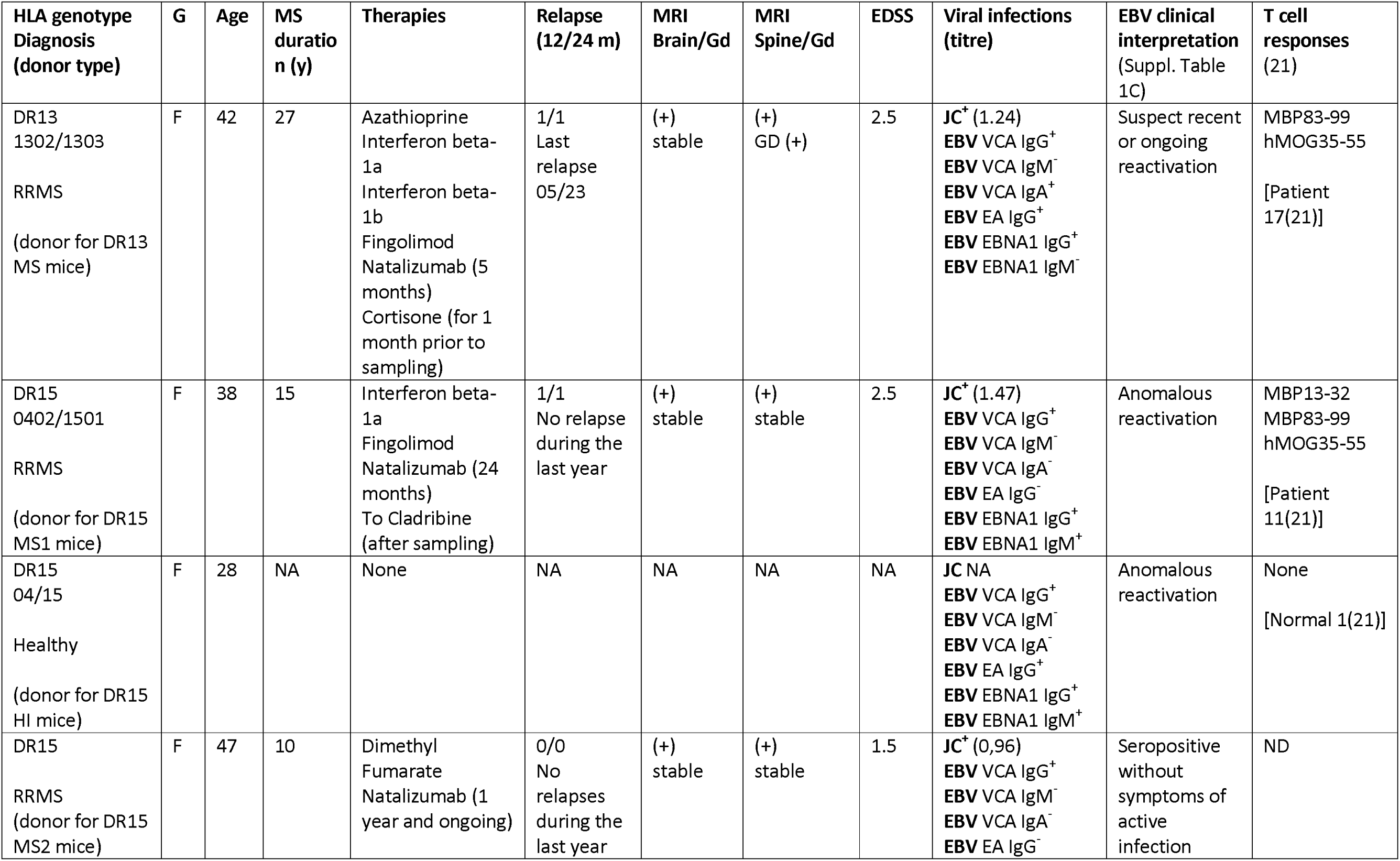
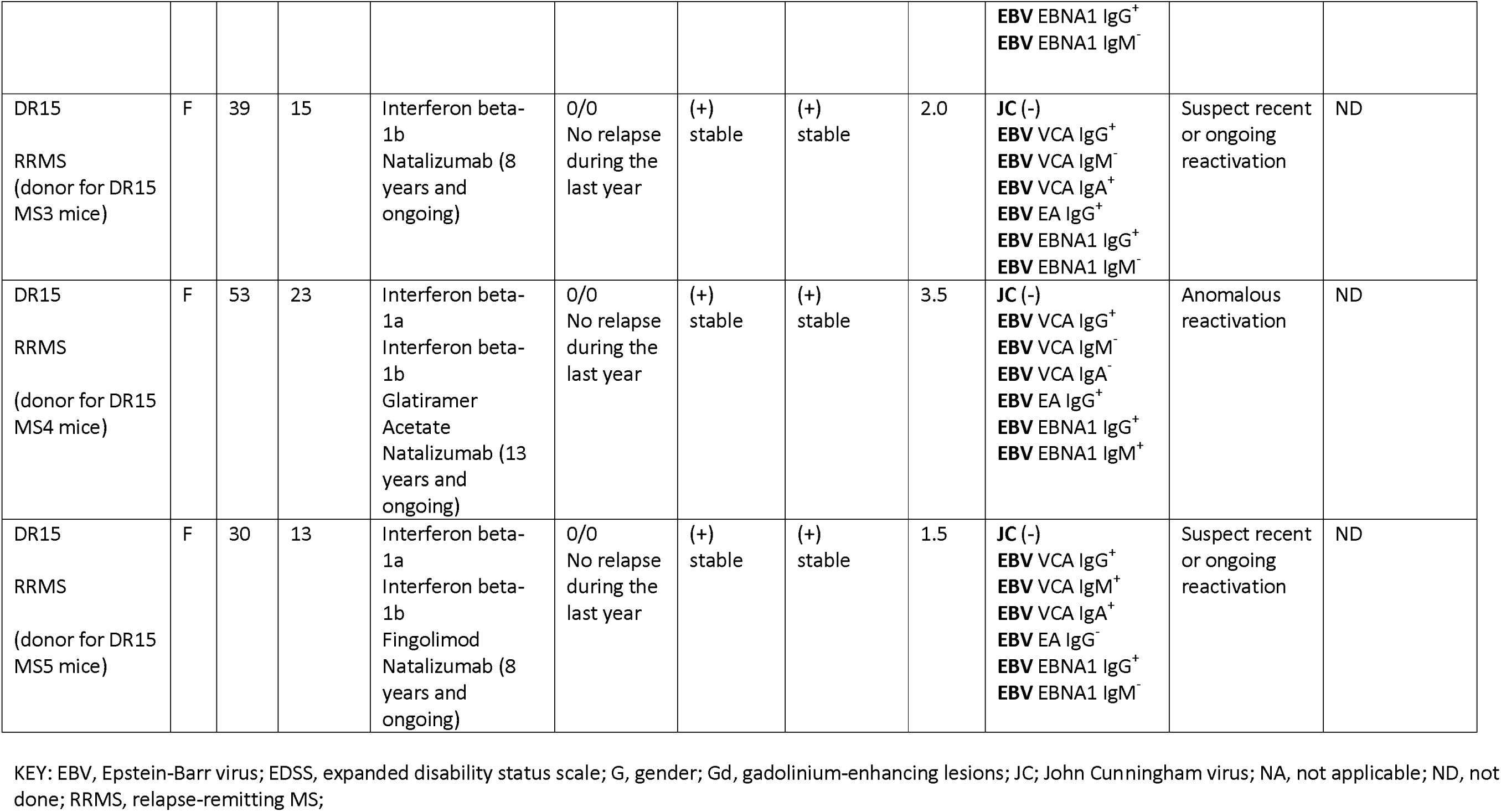

Successful engraftment of mice by hCD45^+^ leukocytes was confirmed in the spleen recovered from non-immunized and immunized mice at sacrifice on dpt 42. Further analysis of splenocytes revealed robust engraftment of hCD4^+^ and hCD8^+^ T cells in all mice, and of hCD19^+^ B cells in DR13 MS, DR15 MS3 and DR15 MS5 mice (Fig. 1C; Supplementary Table 4). Notably, B cell engraftment was best in MS patients interpreted as having suspected recent or ongoing reactivation of EBV, as determined by plasma levels anti-EBV antibodies (Table 1; Supplementary Table 2C). Human immune cells other than T and B cells, notably monocytes, were undetectable in dpt 42 spleen. Analysis of cytokine production by intracellular staining showed high proportions of IFN-γ-producing hCD4^+^ and hCD8^+^ splenocytes, and IL-17A-producing hCD4^+^ and hCD4^-^ splenocytes in both immunized and non-immunized mice (Fig. 1D).

To estimate the contribution of mouse myeloid cell populations in the humanized mice, we analyzed peripheral blood from naïve B2m-NOG mice, and from naïve and CFA-immunized (day 8 post-immunization) NOD-*scid* parental strain and wild-type C57BL/6 (B6) mice, by flow cytometry. Immature Ly6C^hi^ monocytes and Ly6G^+^ cells are known to massively expand in the periphery of B6 mice following immunization with the adjuvants used for EAE, and are critical for the development of EAE (22,23). Unlike B6 mice, very high proportions of mouse (m) CD11b^+^, mCD11b^+^Ly6C^hi^ and mCD11b^+^Ly6G^+^ myeloid cells were already present in the blood of naive B2m-NOG and NOD-*scid* mice, and were not significantly increased by immunization (Supplementary Table 5).

Functionality of the human immune system in B2m-NOG mice was investigated using an *ex vivo* proliferation assay for T cell responses in splenocytes (23), and a cell-based assay for anti-MOG antibody responses in plasma. Briefly, splenocytes recovered from non-immunized and immunized mice at sacrifice were stimulated *ex vivo* individually with the 4 myelin peptides used for immunization (mMOG35-55, hMOG35-55, MOG1-20, MBP83-99). T cell proliferation responses were measured using a CFSE dilution assay. Splenocytes isolated from DR13 MS mice (Fig. 1E), showed significant T cell proliferation responses to all myelin peptides, equal to control splenocytes stimulated with anti-hCD3 antibody. Notably, splenocytes from DR15 MS and DR15 HI mice showed limited or absence of T cell proliferation responses to myelin peptides or polyclonal T cell stimuli, anti-hCD3 antibody and phytohemaglutinin (PHA) (Figure 1-figure supplement 1C). The difference in responses between DR13 and DR15 donors cannot be explained by differences in human B cell engraftment, which are potential antigen-presenting cells in the *ex vivo* T cell proliferation assay, because B cell engraftment was seen in both DR13 MS and DR15 MS mice (Fig. 1C). A previous study showed that T cells isolated from DR15 MS patients display high levels of autoproliferation (24). Therefore, it is possible that poor human T cell proliferation responses in DR15 mice might result from inherently high T cell autoproliferation in the DR15 MS donor PBMC prior to engraftment, although this possibility needs to be formally tested. Anti-hMOG IgG antibody responses were not detectable in the plasma of any mice at any time point.

We next investigated whether draining lymph nodes (LN) structures could be identified in the immunized mice at sacrifice. Mice lacking the common cytokine receptor γ-chain gene are known to have poor lymphoid organ structure and functioning, because NK cells are needed for the formation of LN inducer cells, which in turn are needed for LN organogenesis (25). Nevertheless, the inguinal fat tissue from several immunized mice showed accumulated masses of leukocytes containing mCD45- and hCD45-positive leukocytes, CD8-, CD4- and CD20-positive lymphocytes by immunohistochemistry, although lacking obvious LN organization (Fig. 1F). Together the results show that human T and B lymphocytes are efficiently engrafted in B2m-NOG mice by PBMC from MS patients, that functional T cell responses can be detected depending upon the PBMC donor, while lack of organized LN and germinal centers prevents the generation of anti-MOG IgG antibody responses.

### Human T cells accumulate at CNS borders in non-immunized DR15 MS and DR15 HI mice, and form spontaneous parenchymal lesions in brain and spinal cord of DR15 MS mice

For the period of the study, none of the engrafted mice showed clinical symptoms of neuroinflammation or EAE using criteria commonly used for scoring EAE (21), or other neurological deficits (26). Mild symptoms of GVHD, specifically fur ruffling and reduced mobility (27), were recorded in few mice independent of group close to the time of sacrifice. To determine whether human immune cells enter CNS tissues in PBMC humanized B2m- NOG mice, we performed immunohistochemical analyses of serial sections from brain and spinal cord recovered from non-immunized mice at sacrifice using marker antibodies for human and mouse immune cells as well as for mouse microglia and astrocytes. To further monitor GVHD development we also performed standard histological analyses of lung and liver, two main tissue targets of GVHD.

DR13 MS humanized mice showed very few hCD45-positive leukocytes at CNS borders, specifically meninges, and none in CNS parenchyma, and no further analysis was made (data not shown). DR15 HI mice showed an accumulation of human immune cells at CNS borders, particularly the meninges and some in the choroid plexus of the interventricular foramen between lateral and third ventricles of the brain, and few cells scattered throughout the brain parenchyma (Fig. 2A-C). Both brain border and parenchymal hCD45-positive immune cells comprised mainly hCD8-positive T cells (Fig. 2D), leading to low hCD4/hCD8 T cell ratios, especially in parenchyma (Fig. 2E). Mild activation of Iba1-positive microglia and GFAP-positive astrocytes was seen at sites of immune cell accumulation at CNS borders (Fig. 2C). In spinal cord, hCD3-positive T cells accumulated at borders and few individual cells infiltrated parenchyma of white and grey matter (Fig. 3A-C), forming rare small T cell lesions (≥ 3 adjacent cells) in the white matter (Fig. 3A, bottom left panel, arrowhead). Spinal cord-infiltrating cells in DR15 HI mice also comprised mainly hCD8-positive T cells, leading to low hCD4/hCD8 T cell ratios (Fig. 3D). Mouse CD45-positive cells were rare at barriers, absent from parenchyma, and demyelination was not observed in the brain or spinal cord of DR15 HI mice.

**Figure 2.**
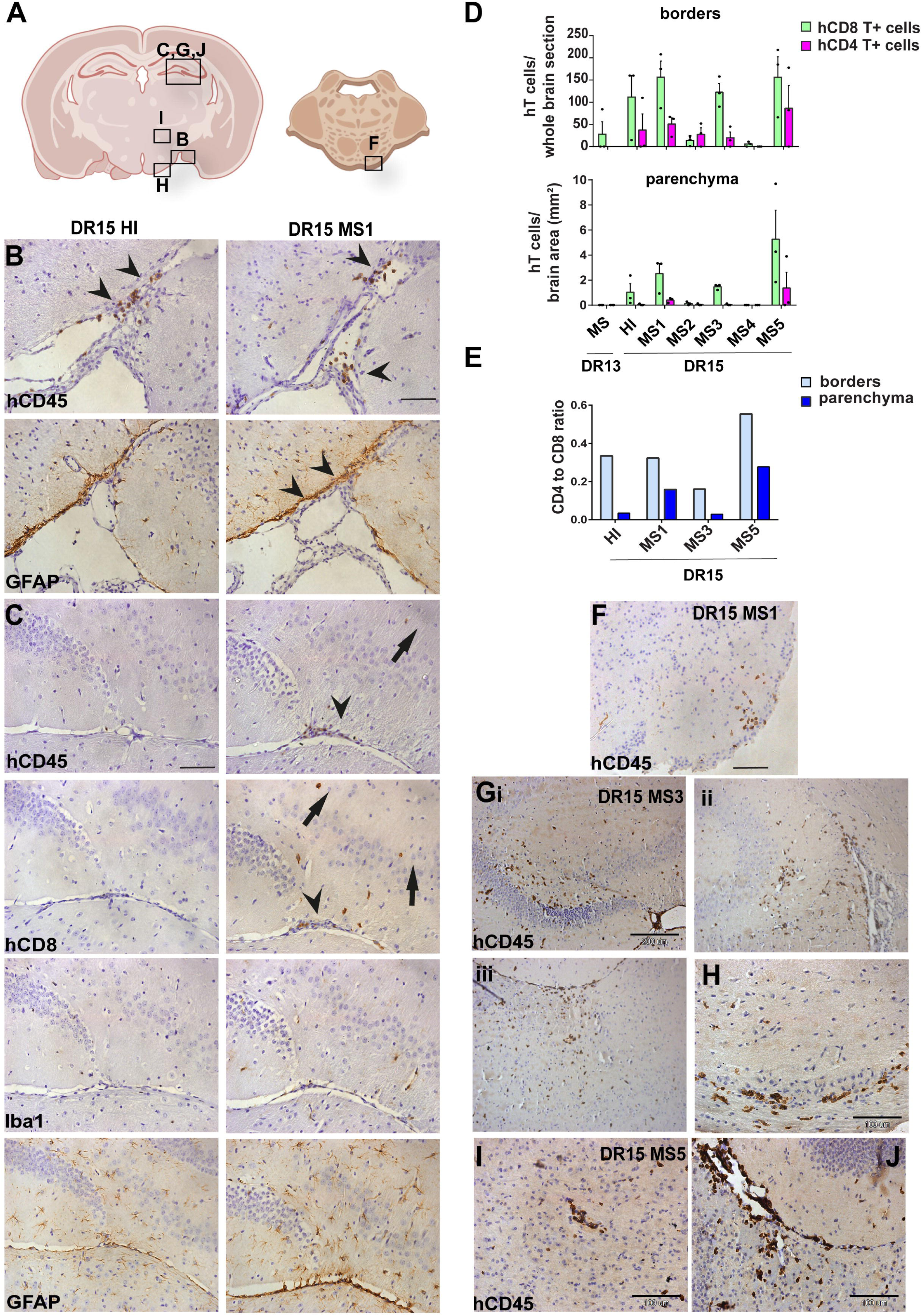
Human T cells accumulate at brain borders in non-immunized DR15 MS and DR15 HI mice, and form spontaneous parenchymal lesions in brain of DR15 MS mice. Immunohistochemical analysis of the brain from non-immunized PBMC B2m-NOG mice showing infiltration by human (h) and mouse (m) CD45-positive leukocytes, and hCD8- positive T cells, in brain border and parenchymal regions (denoted in diagrams of brain and brainstem **A**). **(B, C)** Accumulation of hCD45- and mCD45-positive leukocytes and hCD8- positive T cells, together with local activation of Iba1-positive microglia and GFAP-positive astrocytes, at border regions in the brains of DR15 HI (left panels) and DR15 MS (right panels) mice, specifically at meninges close to the optic tract (B, arrowheads) and in the connective tissue of the interventricular foramen joining the lateral and third ventricles (C, arrowheads). Scattered hCD45-positive leukocytes and hCD8 T cells in brain parenchyma of DR15 MS mice (C, arrows). **(D)** Counting of barrier-associated and parenchymal hCD8- and hCD4-positive T cells at borders and in parenchyma of whole coronal sections of brain of humanized mice, represented as total cells and cells/mm^2^. (**E)** Ratios of hCD4/hCD8 T cells at borders and parenchyma of selected humanized mice. **(F)** Small hCD45-positive immune cell lesion in brainstem of DR15 MS1 mice (see also diagram A). **(G)** hCD45-positive immune cell lesions in grey matter of hippocampus (**i, ii**) and sub-hippocampus/thalamus (**iii**), and white matter of optic chiasm **(H)** of DR15 MS3 mice. **(I, J)** hCD45-positive immune cell lesions in grey matter of thalamus and hippocampus, respectively, of DR15 MS5 mice. Scale bars 100 μm; x20 objective (B, C, F), 200μm; 10x objective (G), 100μm; 40x objective (H, I, J). All results are depicted as mean ± SEM. Statistical analysis was performed by pairwise comparisons between different groups of mice using Student’s t test.

**Figure 3.**
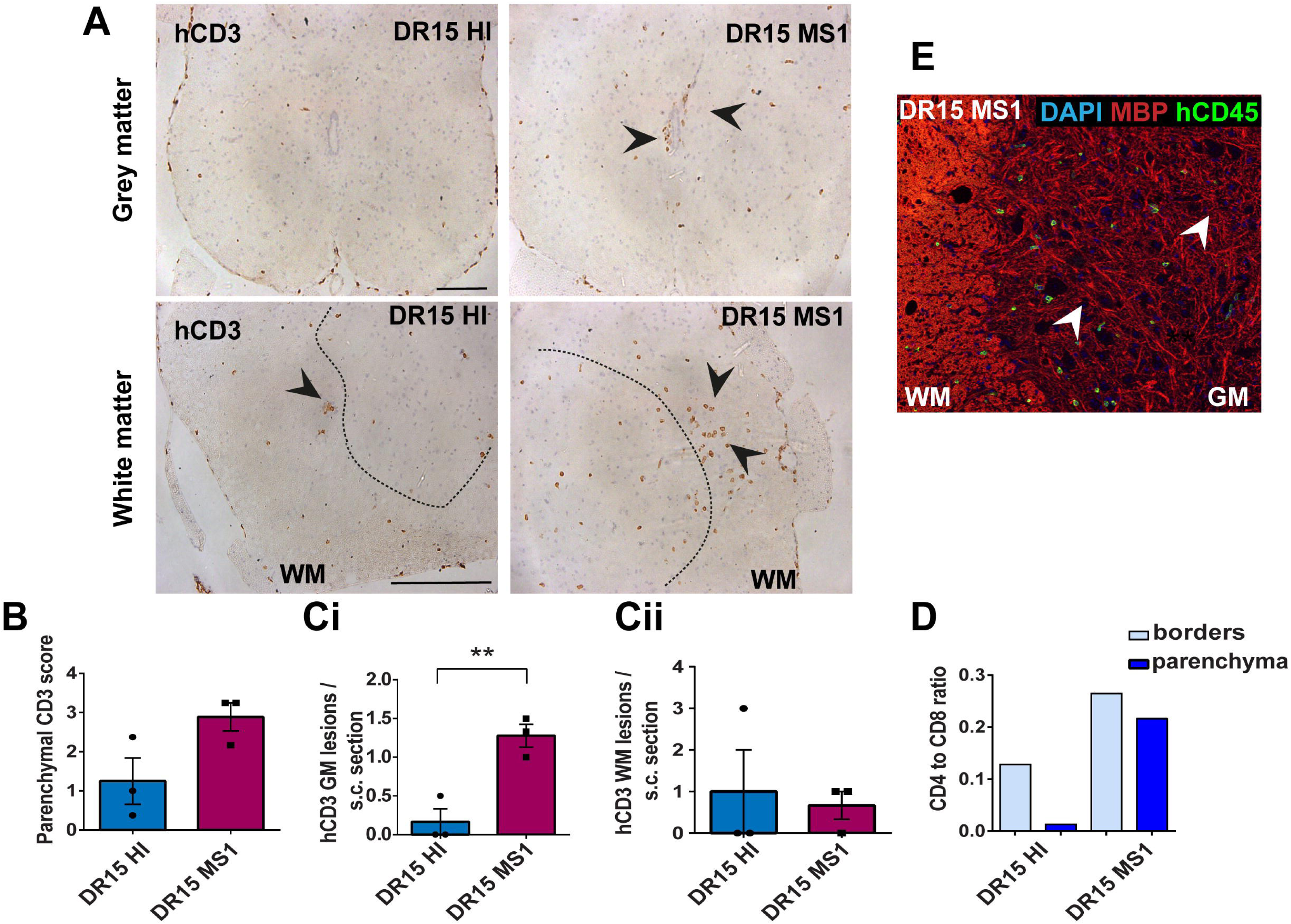
Human T cells infiltrate spinal cord white matter in non-immunized DR15 MS and DR15 HI mice, and form grey matter lesions in DR15 MS mice. Immunohistochemical analysis of spinal cord from non-immunized PBMC B2m-NOG mice showing infiltration by human (h) CD3-positive T cells in the grey and white matter regions. **(A)** Infiltrating hCD3-positive T cells were scattered individually throughout spinal cord grey and white matter and formed small lesions in the white matter (WM) of both DR15 MS and DR15 HI mice (lower panels, arrowheads), and grey matter lesions only in DR15 MS mice (upper panels, arrowheads). The dotted lines mark the boundary between grey matter and white matter. **(B)** Counting of parenchymal hCD3-positive T cell lesions (≥ 3 adjacent cells) in grey and white matter in whole spinal cord sections from DR15 HI and DR15 MS mice. **(C)** Semi-quantitative estimation of hCD3-positive T cells in GM (**i**) and WM (**ii**) regions in whole spinal cord sections compared at equivalent cord regions from DR15 HI and DR15 MS mice. **(D)** Ratios of hCD4/hCD8 T cells at borders and parenchyma of DR15 HI and DR15 MS mice. **(E)** Double immunofluorescence staining for hCD45-positive leukocytes (green, arrowheads) and MBP-positive myelin (red), with DAPI counterstained nuclei (blue), in spinal cord WM of DR15 MS mice. Scale bars 100 μm; x20 objective (A, top panels); x40 objective (A, bottom panels). All results are depicted as mean ± SEM. Statistical analysis was performed by pairwise comparisons between different groups of mice using Student’s t test.

Three of the five groups of DR15 MS mice, specifically DR15 MS1, DR15 MS3 and DR15 MS5 mice, showed severe spontaneous CNS pathology with accumulation of human immune cells at borders, particularly the meninges and choroid plexus of the interventricular foramen between lateral and third ventricles, widespread immune cell infiltration of the parenchyma, and T cell lesions in the brain, brainstem and spinal cord. Brain border and parenchymal hCD45-positive immune cells comprised mainly hCD8-positive T cells (Fig. 2A-D), leading to low hCD4/hCD8 T cell ratios, especially in the parenchyma (Fig. 2E). These mice showed numerous spontaneous parenchymal hCD45-positive cell lesions (≥ 3 adjacent cells). The location and severity of parenchymal lesions were different in each patient, and involved brainstem (DR15 MS1; Fig. 2A, F), hippocampus (DR15 MS3; Fig. 2A, Gi, ii), optic chiasm (DR15 MS3; Fig. 2A, H), and sub-hippocampal regions (DR15 MS3; Fig. 2A, Giii, DR15 MS5; Fig. 2A, I, J). Locally activated Iba1-positive microglia and GFAP-positive astrocytes were observed at sites of immune cell accumulation (Fig. 2C). DR15 MS2 and DR15 MS4 mice showed only few human immune cells at borders and rare parenchymal lesions. In the spinal cord, hCD3-positive T cells accumulated at borders and infiltrated both white and grey matter parenchyma, forming lesions in both areas (Fig. 3A right panels, arrowheads). Numbers of parenchymal hCD3-positive T cells counted in whole spinal cord sections tended to be increased in DR15 MS1 mice compared to control HI mice (Fig. 3B), and numbers of T cell lesions in the grey matter were significantly increased (Fig. 3Ci). Unlike control HI mice, DR15 MS1 mice showed numerous small T cell lesions in the grey matter, often close to the central canal suggesting this is a main site of human T cell entry into the spinal cord grey matter (Fig. 3A, right top panel, arrowheads). Spinal cord-infiltrating immune cells in DR15 MS1 mice also comprised mainly hCD8-positive T cells, leading to low hCD4/hCD8 T cell ratios (Fig. 3D). In contrast to hCD45-positive immune cells that were scattered in spinal cord parenchyma (Fig. 3E), mouse CD45-positive cells were rare at barriers, absent from parenchyma, and demyelination was not observed in the brain or spinal cord of any DR15 MS mice.

To determine whether differences in the severity of CNS immune infiltration in the immunized mice were secondary to levels of GVHD inflammation in the periphery, we performed semi-quantitative analysis of H&E stained liver and lung sections, which are main targets of GVHD responses (27). The mouse groups showed equal levels of inflammation in both liver and lung, revealing a dissociation between peripheral GVHD and CNS inflammation (Figure 2- figure supplement 1). These results show that non-immunized DR15 MS1 humanized mice show increased infiltration of brain and spinal cord by human T cells compared to non-immunized DR15 HI mice, and uniquely develop spontaneous T cell lesions in spinal cord grey matter and brain parenchyma.

### Immunization with myelin peptides increases hCD4 T cell infiltration of CNS parenchyma resulting in mixed hCD4/hCD8 T cell lesions in brain and spinal cord of DR15 MS and DR15 HI mice

To further investigate whether human immune cells have the potential to induce CNS immunopathology in PBMC humanized B2m-NOG mice, we immunized mice for EAE. In a previous study, humanized HLA-DR2b transgenic mice lacking all mouse MHCII genes were found to require high amounts of peptide antigen than B6 mice to induce clinical EAE (21). Here, EAE tests in wild-type B6 mice showed that disease could be induced by immunization with high amounts of peptides with no evidence of increased morbidity or mortality (Supplementary Table 1A, B). A peptide cocktail containing myelin epitopes previously associated with MS, specifically hMOG35-55, MOG 1-20 and MBP83-99 (28–31), plus mMOG35-55, was chosen for immunization in groups of 12-14-week-old humanized B2m- NOG mice on dpt 14. In the first experiment, groups of B2m-NOG mice engrafted with PBMC from DR13 MS, DR15 MS1 and DR15 HI donors received a repeat immunization with 200 μg/peptide spaced 7 days apart, similar to that used in HLA-DR2b transgenic mice. In the second experiment, groups of B2m-NOG mice engrafted with PBMC from DR15 MS2-5 donors received a single immunization with 100 μg/peptide, to reduce the possibility of T cell tolerance (Supplementary Table 1C). None of the PBMC humanized B2m-NOG mice immunized with myelin antigens developed typical clinical symptoms of EAE, a finding consistent with a previous study in NSG mice humanized with PBMC from healthy donors (20).

To investigate whether immunization for EAE increased immune cell infiltration of the brain and spinal cord we performed immunohistochemical analysis of tissues recovered from the mice at sacrifice at dpt 42. Immunized DR13 MS mice and their non-immunized controls showed very few hCD45-positive leukocytes at CNS borders and infiltrating spinal cord parenchyma, so no further analysis was made (data not shown). Immunized DR15 HI mice showed accumulation of hCD45-positive leukocytes at the borders, particularly the meninges and choroid plexus, and few cells scattered throughout the parenchyma (Fig. 4A- C). Brain border and parenchymal human immune cells comprised hCD8-positive T cells, and in contrast to non-immunized controls also contained high numbers of hCD4-positive cells (Fig. 4D), leading to high hCD4/hCD8 T cell ratios (Fig. 4E). This finding is consistent with successful activation of hCD4-positive T cells by the EAE immunization protocol. In one of five immunized DR15 HI mice, a small T cell lesion was detected in the optic tract (Fig. 4A, B, arrowhead). Locally activated Iba1-positive microglia and GFAP-positive astrocytes were present at sites of immune cell accumulation at borders and in optic tract lesions (data not shown). In spinal cord, hCD3-positive T cells accumulated at borders and individual cells infiltrated parenchyma of grey and white matter (Fig. 5A left panels, B), forming small T cell lesions in the white matter (Fig. 5A, bottom left panel, arrowhead, Cii). In the spinal cord of immunized DR15 HI mice, border and parenchymal T cells comprised hCD8-positive T cells and also higher numbers of hCD4-positive T cells compared to non-immunized controls, leading to higher hCD4/hCD8 T cell ratios (Fig. 5D). Mouse CD45-positive cells were rare at barriers, absent from parenchyma, and demyelination was not observed in the brain or spinal cord of immunized DR15 HI mice.

**Figure 4.**
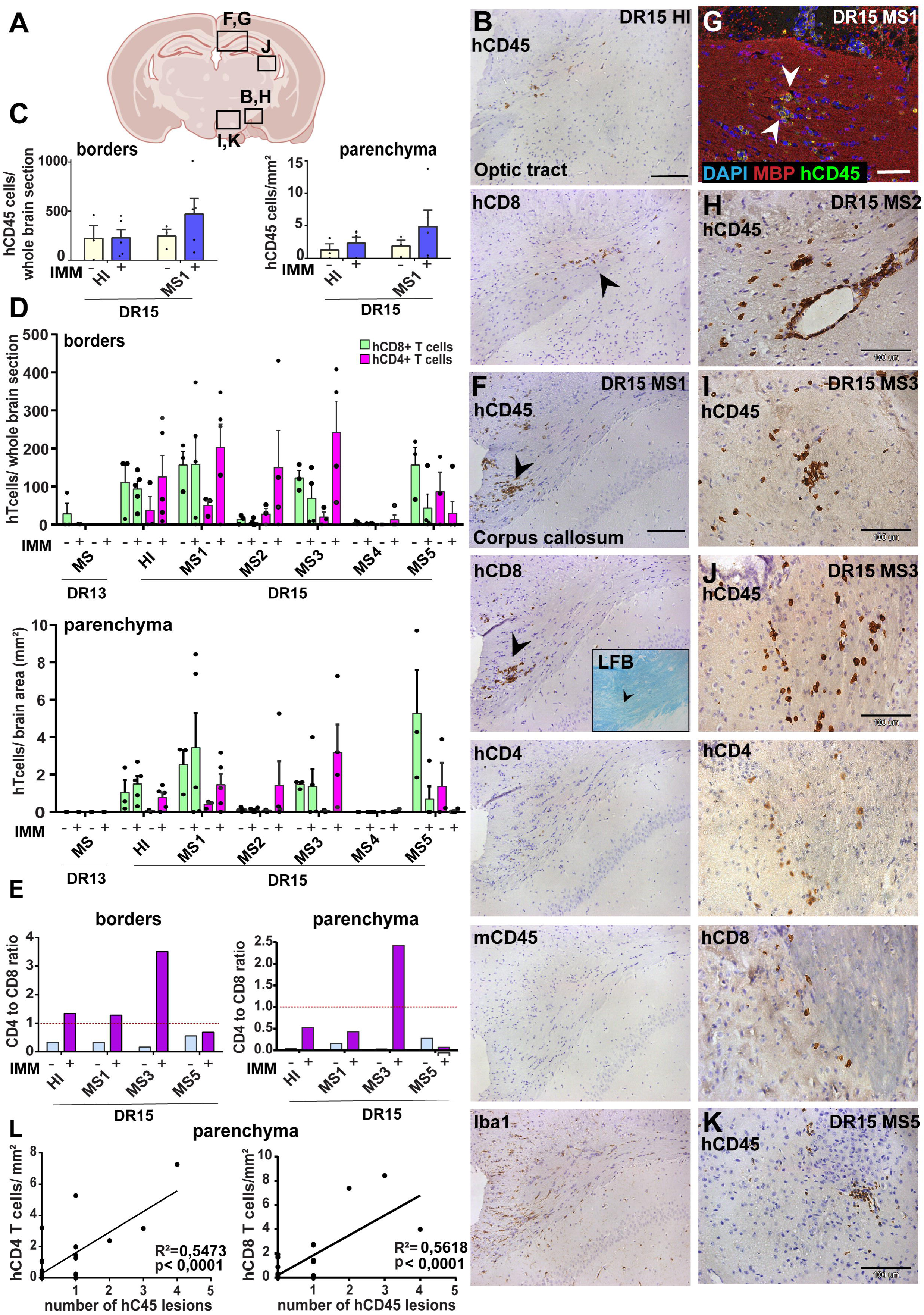
Immunization with myelin peptides increases hCD4 T cell infiltration of brain parenchyma resulting in mixed hCD4/hCD8 T cell lesions in brain of DR15 MS and DR15 HI mice. Immunohistochemical analysis of the brain from PBMC B2m-NOG mice immunized for EAE showing infiltration by human (h) and mouse (m) CD45-positive leukocytes, hCD8- and hCD4-positive T cells, and local activation of Iba1-positive microglia and GFAP-positive astrocytes, in brain regions denoted in the diagram **(A)**. **(B)** Individual hCD45-positive leukocytes and hCD8-positive T cells form small lesions in the optic tract (arrowhead) in immunized DR15 HI mice. **(C)** Counting of hCD45-positive immune cells at borders (total cells in section) and parenchyma (cells/mm^2^) in whole coronal sections of brain from non-immunized and immunized DR15 HI and DR15 MS mice. **(D)** Counting of hCD4- and hCD8- positive T cells at borders (total cells in section) and parenchyma (cells/mm) in whole coronal sections of brain from non-immunized and immunized DR13 MS, DR15 HI and DR15 MS mice. **(E)** Ratios of hCD4/hCD8 T cells at borders and parenchyma of non-immunized and immunized DR15 HI and DR15 MS mice. **(F)** Prominent lesions in the corpus callosum white matter of two of five DR15 MS1 mice (arrowheads), containing hCD45- and mCD45-positive leukocytes, hCD4- and hCD8-positive T cells and locally activated Iba1-positive microglia. Inset shows a serial section stained by Luxol fast blue showing absence of demyelination. **(G)** Double immunofluorescence staining for hCD45-positive leukocytes (green, arrowheads) and MBP-positive myelin (red), with DAPI counterstained nuclei (blue), in corpus callosum in immunized DR15 MS1 mice, showing inflammatory lesion without demyelination. **(H)** Small white matter lesion in DR15 MS2 mouse. **(I)** Prominent white and **(J)** grey matter lesions containing both hCD4 and hCD8 T cells in DR15 MS3 mice. **(K)** Small lesion containing human hCD45-positive immune cells in sub-thalamic area of DR15 MS5 mice. **(L)** Correlation analysis between numbers of hCD4 or hCD8 with number of hCD45 lesions in brain parenchyma of all combined immunized DR15 HI and DR15 MS1-5 mice. Scale bars 100 μm; x20 objective (B, F); x40 objective (G,K). All results are depicted as mean ± SEM. Statistical analysis was performed by pairwise comparisons between different groups of mice using Student’s t test.

**Figure 5.**
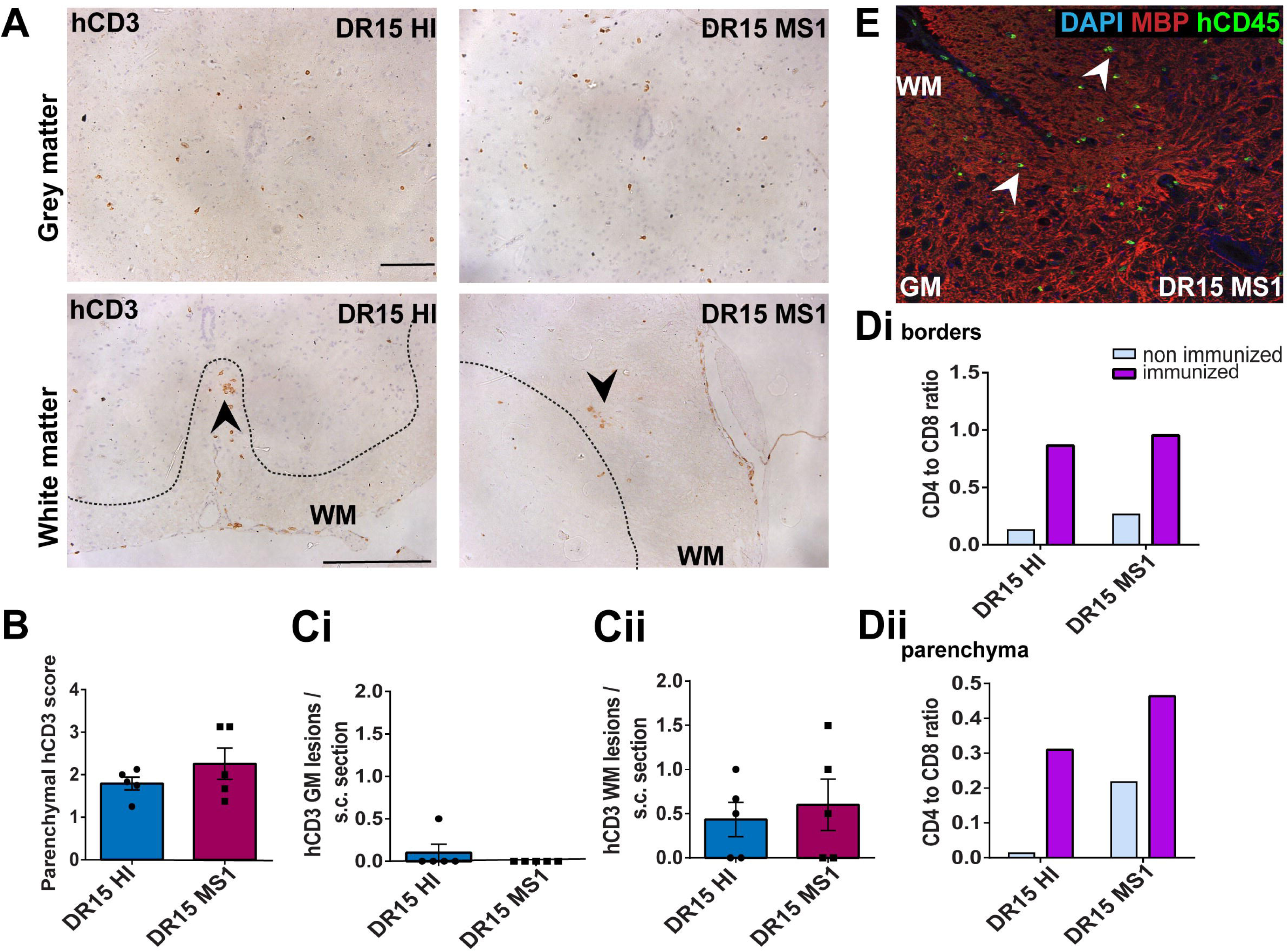
Immunization with myelin peptides increases hCD4 T cell infiltration of spinal cord white matter in both DR15 MS and DR15 HI mice. Immunohistochemical analysis of spinal cord from PBMC B2m-NOG mice immunized for EAE showing infiltration by human (h) CD3-positive T cells in the grey and white matter regions. **(A)** Infiltrating hCD3-positive T cells scattered throughout spinal cord grey and white matter and forming small lesions in the white matter (WM) of both DR15 MS1 and DR15 HI mice (bottom panels, arrowheads) The dotted lines mark the boundary between grey matter and white matter. **(B)** Semi-quantitative estimation of hCD3-positive T cells in whole spinal cord sections from DR15 HI and DR15 MS mice. **(C)** Counting of hCD3-positive T cell lesions (≥ 3 adjacent cells) in grey (GM) **(i)** and white matter (WM) **(ii)** in whole spinal cord sections from DR15 HI and DR15 MS1 mice. **(D)** Ratios of hCD4/hCD8 T cells at borders **(i)** and in parenchyma **(ii)** of spinal cord in non-immunized and immunized DR15 HI and DR15 MS mice. **(E)** Double immunofluorescence staining for hCD45-positive leukocytes (green, arrowheads) and MBP-positive myelin (red), with DAPI counterstained nuclei (blue), in white matter (WM) of immunized DR15 MS spinal cord. All results are depicted as mean ± SEM. Scale bars 100 μm; x40 objective (A). Statistical analysis was performed by pairwise comparisons between different groups of mice using Student’s t test.

Immunized DR15 MS1-5 mice generally showed prominent accumulation of human immune cells at CNS borders, widespread immune cell infiltration of the parenchyma and development of T cell lesions in the brain and the spinal cord. Brain border and parenchymal hCD45-positive leukocytes comprised hCD8-positive T cells together with higher numbers of hCD4-positive cells compared to non-immunized controls (Fig. 4C, D), leading to higher hCD4/hCD8 T cell ratios (Fig. 4E). Border and parenchymal T cells were generally increased in immunized DR15 MS mice compared to immunized DR15 HI mice, but differences did not reach significant due to intra-group variation (Fig. 4D). Unique features of immunized DR15 MS mice included large human immune cell lesions in the corpus callosum white matter of DR15 MS1 mice containing both hCD8- and hCD4-positive T cells (Fig. 4A, F), without detectable myelin damage (Fig. 4F inset, G), a white matter lesion in DR15 MS2 mice (Fig. 4A, H), numerous white and grey matter lesions in DR15 MS3 mice (Fig. 4A, I, J), and small sub thalamic grey matter lesions in DR15 MS5 mice (Fig. 4A, K). Increased brain infiltration by CD4-positive T cells in immunized mice was not associated with increased frequency of brain lesions, but rather the presence of mixed lesions containing both CD4- and CD8- postive T cells (Fig. 4J). Correlation analysis showed a positive correlation between numbers of hCD4 and hCD8 with numbers of hCD45 lesions (Fig. 4L). Locally activated Iba1-positive microglia and GFAP-positive astrocytes were present at sites of human immune cell entry at CNS borders and in T cell lesions (Fig. 4F, and data not shown). In spinal cord, hCD3-positive T cells accumulated at borders and individual cells infiltrated parenchyma of grey and white matter (Fig. 5A right panels, B), forming small T cell lesions only in the white matter (WM) (Fig. 5A, right bottom panel, arrowhead; Cii). Numbers of parenchymal hCD3-positive T cells counted in whole spinal cord sections, and numbers of T cell lesions in the white matter were equal to those in immunized HLA-DR15 HI mice (Fig. 5B). Spinal cord border and parenchymal T cells of immunized DR15 MS1 mice comprised hCD8-positive T cells and also high numbers of hCD4-positive T cells compared to non-immunized controls, leading to higher hCD4/hCD8 T cell ratios (Fig. 5D). Mouse CD45-positive cells were rare at barriers, absent from parenchyma, and demyelination was not observed in the brain (Fig. 4F) or spinal cord of immunized DR15 MS mice.

To determine whether CNS immune infiltration in the immunized mice was secondary to levels of GVHD inflammation in the periphery, we performed semi-quantitative analysis of liver and lung sections with H&E staining. Immunized DR13 MS mice showed the lowest levels of immune cell infiltration in liver and lung, DR15 MS1 mice intermediate levels, and DR15 HI highest levels (Figure 2- figure supplement 1). Results revealed an inverse relationship between peripheral and CNS inflammation in the DR15 HI and MS mice, with DR15 HI mice showing highest peripheral and relatively lower CNS involvement, while DR15 MS mice showed the highest CNS and relatively lower peripheral inflammation. These results show that immunization of DR15 HI and DR15 MS mice with myelin peptides equally induced spinal cord white matter inflammatory lesions, and uniquely induced large confluent brain white and grey matter lesions in DR15 MS mice. Taken together, our results show that PBMC from DR15-positive MS patients show increased propensity to induce inflammatory CNS lesions compared to PBMC from a DR13-positive MS patient or a DR15- positive healthy individual.

## Discussion

Here we show that PBMC from RRMS and healthy donors with different HLA-DRB1 genotypes have the potential to efficiently engraft B2m-NOG mice with human T and B lymphocytes and to induce sub-clinical CNS inflammation dominated by hCD8^+^ T cells in the brain and the spinal cord. Specifically, mice engrafted with PBMC from DR15-positive MS and healthy donors, and not from a DR13-positive MS donor, showed spontaneous brain and spinal cord infiltration by hCD8^+^ T cells, and increased infiltration by hCD4^+^ following immunization for EAE. CNS inflammation was more severe in mice engrafted with PBMC from several DR15-positive MS donors. These mice uniquely developed large human T cell lesions in brain regions including the corpus callosum with or without immunization with myelin peptides using an EAE protocol. Human T cell lesions were associated with locally activated mouse microglia and astrocytes, but notably lacked infiltrating human or mouse monocytes and demyelination, and therefore did not lead to clinical symptoms of EAE or other neurological defects.

PBMC humanized B2m-NOG mice developed several hallmark features of MS that are not readily reproduced in conventional animal models, such as EAE. First, brain immune infiltrates in DR15 MS engrafted mice were enriched for hCD8^+^ over hCD4^+^ T cells, even though the B2m-NOG model is characterized by increased engraftment of CD4^+^ over CD8^+^ T cells (18) (Figure 1B). The preferential infiltration of the CNS parenchyma by CD8^+^ T cells and the presence of inflammatory lesions in the brain white and grey matter in our model are both features of MS (3,8,32). Second, T cell lesions and locally activated microglia and astrocytes, were observed in brain white matter regions, particularly in the corpus callosum and optic tracts. Lesions in the corpus callosum are important findings in MRI of MS patients and are correlated with cognitive problems as well as disability in the upper limbs (33). Inflammation in the optic tracts is often a first presenting symptom in MS and related disorders (34). Third, human leukocytes were concentrated on the meninges and choroid plexus, notably in the interventricular foramen connecting the lateral and third ventricles. The choroid plexus is an important immunological site for immune surveillance and a site of CD8^+^ T cell and granulocyte involvement in progressive MS (35).

Nevertheless, MS is a complex disease and immunopathology was only partially reproduced in the brain and the spinal cord of PBMC humanized B2m-NOG mice. Parenchymal lesions contained both hCD4^+^ and hCD8^+^ T cells associated with locally activated microglia and astrocytes, but there was poor engraftment by human monocytes in the blood, spleens or CNS of both immunized and non-immunized B2m-NOG mice. Blood myeloid cells are short-lived and constantly renewed by the bone marrow, so that human myeloid cells are nearly absent in the commonly used PBMC-NSG model (18). In contrast, mouse CD45^+^ cells comprising very high proportions of CD11b^+^ myeloid cells, Ly6C^hi^ monocytes and Ly6G^+^ cells, were present in the blood of immunized and non-immunized NOD-*scid* mice, which are a parental strain for B2m-NOG mice, and the non-immunized B2m-NOG mice prior to engraftment. Nevertheless, mouse CD45^+^ myeloid cells did not infiltrate the CNS of the humanized mice, even after EAE induction. The absence of CNS-infiltrating human or mouse monocytes in the PBMC humanized B2m-NOG mice NOG mice is most likely to be responsible for the absence of clinical symptoms and demyelination in this system. Our results are consistent with a previous study in which NSG mice were humanized with PBMC from healthy donors and immunized for EAE (20). In that study, mice developed CNS inflammation with infiltrating CD4^+^ and CD8^+^ T cells but did not show clinical symptoms or demyelination (20). CNS-infiltrating Ly6C^hi^CCR2^+^ monocytes, which differentiate into inflammatory macrophages locally in the CNS after being activated by T cells, are critical effector cells in EAE (22,36). These cells are massively recruited into the periphery from the bone marrow following immunization (23), and are likely to compete for space with engrafted human immune cells before and after EAE induction. The use of immunodeficient NOG/NSG models expressing human transgenes that would support engraftment of human monocytes, or facilitate communication of human T cells with mouse inflammatory monocytes (37,38), should allow a more comprehensive modeling of MS immunopathology in humanized mice.

Another limitation for MS modeling in the B2m-NOG mice used in this study is the poor LN structure and functioning that results from depletion of the common cytokine receptor γ-chain gene (25). This deficiency was confirmed in this study by the absence of organized LN and germinal follicles, and of anti-MOG IgG antibodies in the PBMC humanized B2m-NOG mice immunized for EAE. Again, further development of available immunodeficient mouse strains to promote LN organogenesis will be an important step for modeling MS immunopathology in mice. In a previous study, NSG mice injected directly in the CNS with CSF cells from MS patients (39) developed clinical symptoms, immune infiltration of the CNS and demyelination, although differences in the route of administration of the transplanted cells might explain the differences between the results of that study and those presented here and by others (20).

Our observation that mice engrafted with PBMC from DR15-positive active RRMS patients showed both spontaneous and EAE-inducible T cell lesions in the brainstem and spinal cord is highly relevant in the context of previous data concerning this haplotype. HLA-DR2 has long been associated with MS (40), with the DR15 haplotype being linked to earlier disease onset (1,41). MHCII molecules present antigens to CD4^+^ T cells, and DR15 is strongly associated with the autoproliferation of peripheral CD4^+^ T cells in MS patients (42). T cell autoproliferation is mediated by memory B cells in a HLA-DR-dependent manner, and autoproliferating T cells are increased in MS patients treated with natalizumab and enriched for brain-homing T cells (24). In a recent study with humanized NSG mice, CD34^+^ hematopoietic progenitor cells (HSC) from DR15-positive donors induced higher peripheral T cell responses and alloreactivity compared to HSC from DR15-negative donors, and increased CD8^+^ T cell responses after EBV infection (43). In our study human T cells isolated from DR13 MS mice showed positive T cell proliferation responses to myelin peptides and hCD3 antibody. This provides proof-of-principle for successful engraftment of functional human T cells in humanized mice. However, none of the T cells isolated from DR15 MS1-5 mice showed further proliferation ex vivo, either to myelin peptides or to polyclonal stimuli hCD3 antibody and PHA, possibly due to inherent high levels of autoproliferation in these cells (24). Alternatively, CD4+ T cells may become nonresponsive to anti-CD3 antibody once engrafted into B2m-NOG mice, as previously reported in NOD-scid mice (44). Noteworthy is that MS patients suspected of recent or ongoing reactivation of EBV (DR13 MS, DR15 MS3 and DR15 MS5; Table 1), showed the best engraftment of hCD19^+^ B lymphocytes in mice, and that both DR15 MS3 and DR15 MS5 mice showed multiple CNS lesions.

There is now strong evidence that EBV infection plays a major role in MS pathogenesis (45), acting synergistically with DR15 to increase the likelihood of developing MS (40). DR15 individuals with specific antibody reactivity to EBNA1 (amino acid 385–420) have 24-fold increased risk for MS (46). Anti-EBNA-1 antibodies present high homology and react with several CNS antigens, such as αB-crystallin, MBP, anoctamin 2 and more recently glial cell adhesion molecule (47–50). Nevertheless, positive correlations between the DR15 haplotype, T cell autoproliferation, EBV and the development of brain lesions in humanized mice remain to be formally tested in a larger study using greater numbers of DR15-positive MS donors compared to DR15 negative MS and DR15-positive healthy controls, together with the measurement of basal levels of T cell proliferation in the peripheral blood of the PBMC donors.

In summary, PBMC humanized B2m-NOG mice show partial representation of MS immunopathology that is not reproduced in conventional animal models such as EAE, particularly the development of hCD8^+^ lesions in the grey and white matter of the brain and spinal cord. New immunodeficient mouse strains developed to promote human monocyte engraftment and the development of organized LN therefore hold great promise for comprehensive modelling of MS immunopathology. An important outcome of this study is that PBMC humanized B2m-NOG mice highlight the variability between the different human PBMC donors, and therefore represent a simple and rapid approach for generating personalized immune models for drug screening.

## Materials & Methods

### MS patients and healthy subjects

Blood donors were selected from a Hellenic MS patient cohort attending the MS outpatient clinic at Aeginition University Hospital of the National and Kapodistrian University of Athens (NKUA) School of Medicine, supervised by Dr. Maria Anagnostouli and genotyped for the HLA-DRB1 allele, at the Research Immunogenetics Laboratory, of the same Hospital, as previously described (21) (Table 1). Six patients with long-term RRMS recently presenting with highly active disease since the last dose of immunomodulatory therapy (natalizumab) were selected. Natalizumab treatment was chosen as a criterion for this study because it blocks lymphocyte migration to the brain, thereby increasing circulating T and B lymphocytes (51). One patient was DR13-positive (DR13 MS), and five were DR15-positive (DR15 MS). The DR13-positive patient additionally received cortisone treatment 1 month prior to sampling because of high disease activity. A healthy DR15-matched healthy individual (DR15 HI) was selected as control. Donor characteristics, including viral infections and EBV status are listed in Table 1. Blood samples were obtained from MS patients and a healthy individual under signed informed consent in accordance with the Declaration of Helsinki and approval from the Institutional Ethics Committee of Aeginition Hospital, NKUA (Protocol No: 7BSH46Y8N2-B66, 13/05/2015).

### PBMC isolation

Fresh peripheral blood (50 ml) was collected in EDTA-treated polypropylene tubes and PBMC were isolated under Ficoll-Histopaque®-1077 gradient centrifugation (Sigma-Aldrich). Buffy coats containing blood leukocytes were washed with 2% FBS in PBS. Erythrocytes were removed using Quicklysis™ erythrocyte lysis buffer (Cytognos) and leukocytes were washed with 2% FBS in PBS. Before transfer of PBMC into mice, we compared several protocols for preparation of human PBMC using blood from a healthy individual, specifically, 1) PBMC isolated from fresh blood on the day of transfer, 2) PBMC isolated from blood after 3 days’ storage at 4°C, 3) PBMC cultured for 24 h in complete RPMI-1640 after snap freezing of fresh PBMC and thawing. Cell viability and analysis of immune cell populations by flow cytometry showed that freshly isolated PBMC gave the best results and were used for transfer in this study (Supplementary Table 3, protocol 1).

### EBV antibody responses in human blood

Blood plasma from the DR13 MS, DR15 MS and DR15 HI donors was analyzed for EBV antibody responses using the following ELISA tests: ELISA-VIDITESTS anti-VCA EBV IgG (ODZ-265), anti-VCA IgM (ODZ-005), and anti-VCA IgA (ODZ-096), anti-EA (D) EBV IgG (ODZ-006), anti-EBNA-1 EBV IgM (ODZ-002) and anti-EBNA-1 EBV IgG (ODZ-001) (Vidia). Standardized criteria for the clinical interpretation of EBV antibody responses are shown in Supplementary Table 2C.

### Mice

Ten-to-twelve-week-old female immunodeficient B2m-NOD/Shi - *scid IL2rg*^null^ (B2m-NOG) mice (stock number 14957-F: NOD.Cg-B2m<em1TAC> Prkdc <scid>Il2rg<tm1Sug>JicTac; Taconic Biosciences) were used for transfer of freshly isolated human PBMC. These mice lack mature T, B, and NK cells and lack MHC class I molecules. Previous studies show that an equivalent mouse strain, *NOD.Cg-Prkdc*^scid^*Il2rg*^tm1Wjl^*B2m*^tm1Unc^*/Sz* (NSG β2m^null^; The Jackson laboratory) facilitates the engraftment of CD4^+^ over CD8^+^ T cells and exhibits delayed onset of graft versus host disease (17,18). NOD.CB17-Prkdc scid/NCrHsd (Envigo) mice were used for investigation of peripheral mouse myeloid cell populations in a related severely immunodeficient mouse strain by flow cytometry.

Mice were transplanted via tail vein injection with 10x10^6^ PBMC in 200 μL PBS on the same day as cell isolation from fresh peripheral human blood. Mice were assessed daily for symptoms of GVHD, such as fur ruffling, hunched posture, reduced mobility and tachypnea (27), as well as for any spontaneous neurological deficits (26).

All experiments were performed under sterile conditions in the Department of Animal Models for Biomedical Research of the Hellenic Pasteur Institute. Mice were housed in a specific pathogen-free facility in microisolator cages and given *ad libitum* UV-sterilized standard chow, acidified water and gels for hydration. The experiments complied with ARRIVE guidelines and were in accordance with the local Ethical Committee guidelines on the use of experimental animals at the Hellenic Pasteur Institute, were approved by the national authorities and complied to EU Directive 2010/63/EU for animal experiments. The animal study was reviewed and approved by Committee for Evaluation of Experimental Procedures, Department of Experimental Animal Models, Hellenic Pasteur Institute (Presided by Dr P Andriopoulos the Hellenic Republic, General Secretariat for Agricultural Economy, Veterinary and Licenses), and performed under licence numbers 770851/27-11-2019 and 1343917/02-11-2023.

### Immunization with myelin peptides

Peptides MBP83-99, MOG1-20, and mMOG35-55 and hMOG35-55 (S42 in mMOG, P42 in hMOG), were synthesized as previously described (52). In a previous study, we found that humanized HLA-DR2b transgenic mice lacking all mouse MHCII genes required greater amounts of peptide antigen to induce clinical EAE than wild-type B6 mice (21). Due to limited numbers of B2m-NOG mice available for this study, several EAE protocols were first tested using wild-type B6 mice, mainly to exclude the possibility of toxicity at high peptide doses. Briefly, 8-week-old female B6 mice were immunized by subcutaneous (s.c.) tail-base injection of 37 μg (our standard EAE protocol) or 200 μg mMOG35-55, or a myelin peptide antigen cocktail containing 200 μg each of mMOG35-55, hMOG35-55, MOG1-20 and MBP83-99, dissolved in 100 μl saline and emulsified in an equal volume of Freund’s complete adjuvant (FCA) (Sigma-Aldrich). FCA was supplemented with 400 μg/injection of H37Ra *Mycobacterium tuberculosis* (Difco, BD Biosciences). All mice, except those immunized using the standard protocol, received an identical boost immunization 7 days later. The immunized mice also received intraperitoneal (i.p.) injections of 200 ng of *Bordetella pertussis* toxin (PTx) (Sigma-Aldrich) at the time of each immunization and 48 h later. All mice developed clinical symptoms of EAE with no evidence of increased morbidity or mortality at high peptide doses. For this reason, the high-dose peptide cocktail was chosen for immunization of groups of 12-14-week-old PBMC humanized B2m-NOG mice on dpt 14 in EAE experiment 1. In EAE experiment 2, groups of 12-14-week-old PBMC humanized B2m-NOG were immunized once, using 100 μg each peptide/mouse, to reduce the possibility of immune tolerance. Mice were monitored daily for the clinical symptoms of EAE according to criteria commonly used for scoring EAE: 0, normal; 1, limp tail; 2, hind limb weakness; 3, hind limb paralysis; 4, forelimb paralysis; 5, moribund or dead (0.5 gradations represent intermediate scores) (21). Mice were also monitored for signs of other neurological deficits (26).

### MOG antibody responses in PBMC humanized mouse blood

Peripheral blood was collected from immunized and non-immunized mice at sacrifice on dpt 42, and sera collected for analysis of antibody responses. Samples were screened for IgG antibodies against MOG, using indirect immunofluorescence assays (IFA). Sera were analyzed at a starting dilution of 1:10, on EU 90 cells transfected with MOG protein and on EU 90 control transfected cells, according to the manufacturer instructions (Euroimmun, Lubeck, Germany). Visualization and evaluation of IFA results were performed under a fluorescence microscope (Zeiss, Axioskop 40).

### Flow cytometry

Fresh peripheral blood samples from the MS and HI donors were analyzed by FACS using a panel of immune cell marker antibodies. To monitor immune cell populations, 3 ml fresh peripheral blood was collected in EDTA tubes and a 100 μl aliquot was used for flow cytometry. Erythrocyte lysis was achieved with with Quicklysis^TM^ (Cytognos) for 25 min. Cells were washed and incubated with an antibody mixture for human immune cell markers BV- 510-hCD45 (clone 2D1), APC-Cyanine 7-hCD3 (clone HIT3a), PE-Cyanine 7-hCD4 (clone A161A1), PerCP-hCD8 (clone SK1), PE-hCD19 (clone 4G7), FITC-hCD56 (clone 5.1H11), APC- hCD66b (clone G1OF5), BV-421-hCD14 (clone HCD14) (all from Biolegend), for 30 min at 4 °C (Supplementary Table 2A). To monitor engraftment of human immune cells in B2m-NOG mice, peripheral blood was collected in heparin tubes from the tail vein at dpt 7, 13 and 42. Erythrocyte lysis and incubation with antibody mixture was performed as for fresh human blood above, except that the APC-hCD66b antibody was replaced by mouse APC-CD45 (clone 30-F11) (Biolegend) (Supplementary Table 2A). Data was acquired with FACSCelesta^TM^, FACSCanto^TM^ II or FACSMelody^TM^ cytometer and analyzed with FACSDiva™ (BD) and FlowJo software (Tree Star, Inc.).

For intracellular staining of IFN-γ, splenocytes were isolated at sacrifice at dpt 42 as described above, and cells were stimulated with phorbol 12-myristate 13 acetate (PMA) (20 ng/ml, Sigma-Aldrich) and ionomycin (1μg/ml, Sigma-Aldrich) and treated with brefeldin-A (5μg/ml, Sigma-Aldrich) for 3 h at 37°C/5% CO2 in order to disrupt Golgi-mediated transport. Cells were fixed with 2% PFA in FBS, permeabilized using 0.5% wt/vol saponin, and stained with PerCP-Cyanine 5.5-CD3 (clone HIT3a, Biolegend), antibodies for APC-CD4 (clone RPA-T4, BD Biosciences), PerCP-CD8 (clone SK1, Biolegend), and PE-IFN-γ (clone B27; BD Biosciences) and PE-IL-17A (clone eBio64DEC17, eBioscience). Data acquisition was performed using a FACSCalibur^TM^ cytometer and analyzed with FlowJo software (Tree Star, Inc.).

### T cell proliferation assay

Spleens were isolated at sacrifice and cells mechanically separated in RPMI (Invitrogen Life Technologies) containing 10% heat-inactivated FCS. To obtain single cell suspensions, the washed homogenate was passed through a 70 μm cell strainer and erythrocyte lysis was performed with Gey’s erythrocyte lysis buffer for 5 min. The reaction was stopped with RPMI 1640/FCS. Washed splenocytes at a concentration of 10^7^ cells/mL in PBS were incubated with 5 μΜ carboxylfluorescein succinimidyl ester (CFSE, V12883, Thermofisher) for 15 min at 37°C. Cells were washed and PBS/2% FCS was applied for 30 min at 37°C to stop the reaction. Cells were washed and resuspended in RPMI 1640/FCS with 50 μM 2- mercaptoethanol (Sigma-Aldrich) at concentration of 10^6^ cells/ml. The cells were stimulated in triplicates in round-bottom 96-well plates with a myelin peptide antigen cocktail containing 30 μg/ml each mMOG35-55, hMOG35-55, MOG1-20 and MBP83-99 for 120 h. Negative control cells were incubated with medium only, and positive control cells were incubated in plates coated with anti-CD3 (1μg/ml) (clone HIT3a, BD Biosciences) or PHA (2 μg/ml) (Sigma-Aldrich). Results were expressed as cell division index which is the ratio of %CFSE^low^ splenocytes (proliferating splenocytes) cultured with peptides or anti-CD3 to the of %CFSE^low^ splenocytes cultured with medium only. Data acquisition was performed using a FACSCalibur^TM^ cytometer and analyzed with FlowJo software (Tree Star, Inc.)

### Histopathology and Immunohistochemistry

Mice were transcardially perfused with ice-cold PBS at sacrifice on dpt 42 by carbon dioxide inhalation. Brains, spinal cords and peripheral tissues (including lung, liver and inguinal fat tissue) were dissected and post-fixed in 4% paraformaldehyde fixative (PFA) overnight at 4°C. Tissues were embedded in paraffin and 5μm sections used for immunohistochemistry and histopathology. Inflammation in peripheral tissues was visualized by haematoxylin and eosin (H&E) using standard techniques. Immune cell infiltration and inflammation of brain and spinal cord were visualized by immunohistochemistry using antibodies specific for hCD45, hCD4, hCD8, mCD45, and mouse Iba1 and GFAP followed by biotinylated secondary Abs, horseradish peroxidase-labeled avidin-biotin complex, and signal development using 3131-diaminobenzidine. Nuclei were counterstained with hematoxylin. Ιmages were captured with an Olympus DP71 microscope digital camera using cell^A imaging software (Soft Imaging System GmbH). High resolution immunofluorescence imaging was performed by confocal microscopy using paraffin sections immunostained with marker antibodies for myelin (MBP), microglia (Iba1) followed by anti-rat CF-647- (Biotrium) and anti-rabbit AlexaFluor 568- (Invitrogen) labelled secondary antibodies, respectively. Nuclei were counterstained with DAPI. A Leica TCS SP8 confocal microscope was used to acquire fluorescent images.

### Statistical Analysis

Data were processed using Microsoft Excel, and statistical analysis was performed with Graphpad Prism 8. Figures were made using Adobe Illustrator V24.3 Data was analyzed with Student’s t test and results are presented as means ± SEM. Results were considered statistically significant when p ≤ 0.05. Graphical abstract was created using BioRender.com

## Data availability statement

The raw data supporting the conclusions of this manuscript will be made available by the authors, without undue reservation, to any qualified researcher.

## Ethics approvals

The studies involving human participants were reviewed and approved by the Scientific Committee of Aeginition Hospital, National and Kapodistrian University of Athens, Greece. Licence 283/13-05-2015ΑDΑ: 7ΒΣΗ46Y8Ν2-Β66. The patients/participants provided their written informed consent to participate in this study. The animal study was reviewed and approved by Committee for Evaluation of Experimental Procedures, Department of Experimental Animal Models, Hellenic Pasteur Institute (Presided by Dr P Andriopoulos the Hellenic Republic, General Secretariat for Agricultural Economy, Veterinary and Licenses). Licence numbers 770851/27-11-2019 and 1343917/02-11-2023.

## Patient consent statement

Written informed consent was obtained from the human individual(s) for the publication of any potentially identifiable images or data included in this article.

## Authors contributions

IP designed, set-up and performed experimental protocols for PBMC preparation and humanization of B2m-NOG mice, monitored mice for clinical symptoms and performed immunohistochemical analysis of CNS tissues, analysed results and prepared figures; MK performed quantitative immunohistochemical analysis of CNS tissues, analysed results, monitored EAE mice for clinical symptoms and prepared figures; AD designed and performed EAE experiments, monitored mice for clinical symptoms, performed immunohistochemistry of LN tissues and FACS analyses of humanized mice, analysed results and prepared figures; VG performed FACS analysis and EBV ELISA on fresh human donor blood samples; LD designed human FACS antibody panel, performed analyses of human and mouse blood and analysed results; NM collected and co-ordinated delivery of human donor blood samples; FB performed histological analysis of peripheral humanized mouse tissues and evaluated GVHD; TT designed, synthesized, purified and characterized myelin peptide analogues; MA performed HLA genotyping, selected patients for this study, and supervised Hellenic MS patient cohort; LP was responsible for overall experimental design and management of project, analysed and interpreted data, wrote paper. All authors contributed to the article, edited, and approved the submitted version.

## Acknowledgements

We wish to thank Ivan Gladwyn-Ng and Dimitri Gimnopoulos (Taconic Biosciences) for providing B2m-NOG mice and expert scientific support, Konstantinos Kambas (Molecular Genetics Lab, HPI), Maritsa Margaroni (Flow Cytometry Unit, HPI), Katerina Nanou and George Kollias (BSRC Alexander Fleming) for support with the flow cytometry analyses, Maria Belimezi and Manolis Angelakis (Diagnosis Dept, HPI) for screening mouse plasma for anti-hMOG IgG antibodies, and Maria Avloniti for contributing research material. We especially thank Eirini Fragkiadaki and Ariadne Karles, and other members of the Department of Experimental Animals for Biomedical Research of HPI for their help and expert support in maintenance of the B2m-NOG mice, and the training of experimenters in mouse familiarization techniques during these experiments.

---

**Supplementary Figure 1: Reconstitution of a human adaptive immune system in B2m-NOG mice engrafted with PBMC from MS patients. (A)** Progressive engraftment of human (h) CD45+ leukocytes from donors in groups of B2m-NOG mice, non-immunized (left) and immunized (right) for EAE using the single immunization protocol with 100 μg/each myelin peptide (EAE experiment 2), measured in peripheral blood samples taken at different time points, and in spleen recovered at sacrifice 42 days’ post-transplantation (dpt 42). **(B)** Proportions of hCD4^+^ and hCD8^+^ T cells in blood hCD45^+^ CD3^+^ T cells at different time points. **(C)** Proportions of CFSE^low^ (proliferating) splenocytes recovered from non-immunized (-) and EAE-immunized (+) PBMC B2m-NOG mice at sacrifice on day post transplantation 42 (dpt) and cultured for 120h in the presence of individual myelin peptides, anti-CD3 and PHA, or unstimulated (US). All results are depicted as mean ± SEM.

**Supplementary Figure 2: Inflammation in peripheral GVHD target tissues in PBMC B2m-NOG mice.** Histochemical analysis of peripheral tissues recovered from non-immunized and EAE-immunized PBMC B2m-NOG mice at sacrifice on day post-transplantation 42 (dpt 42). Inflammation was evaluated in paraffin tissue sections of lung **(A)** and liver **(Bi)** and stained by haematoxylin and eosin (H&E). Representative photomicrographs of liver from the different mouse groups are shown **(Bii)**.

