## Supplementary figures and images for "Spontaneous human CD8 T cell and EAE-inducible human CD4/CD8 T cell lesions in the brain and spinal cord of HLA-DRB1*15-positive multiple sclerosis PBMC humanized mice"

### Supplementary figure 1

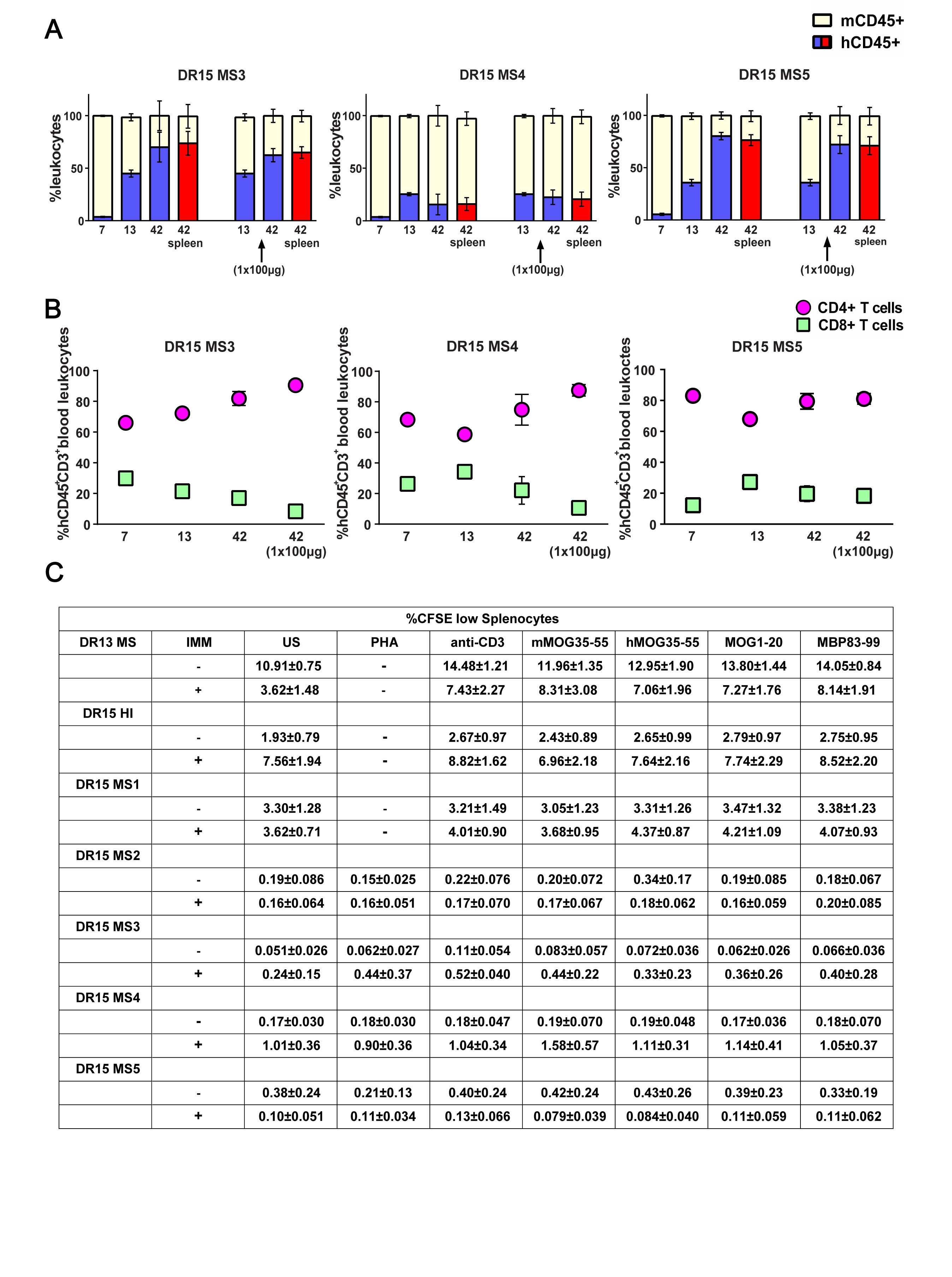

### Supplementary Figure 2

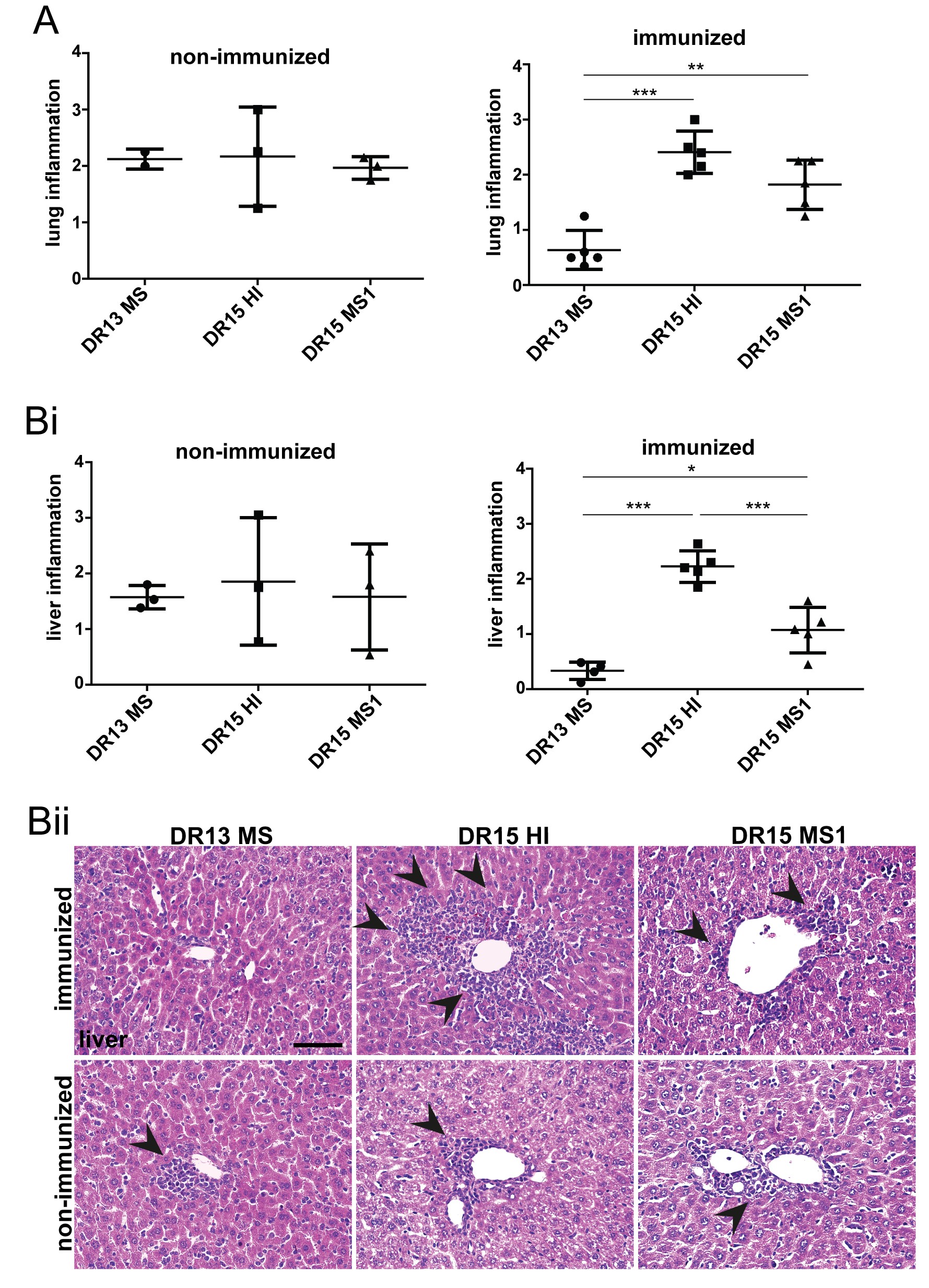
