## Supplementary Table 1 for "Spontaneous human CD8 T cell and EAE-inducible human CD4/CD8 T cell lesions in the brain and spinal cord of HLA-DRB1*15-positive multiple sclerosis PBMC humanized mice"

**Supplementary table 1: EAE protocols used in C57BL/6 and PBMC B2m-NOG mice.** Previously we found that humanized HLA-DR2b transgenic mice lacking all mouse MHCII genes required greater amounts of peptide antigen to induce clinical EAE than wild-type C57BL/6 (B6) mice (21). (**A, B**) To test for possible toxicity of EAE induced by high amounts of myelin peptide antigens, we compared increasing dosages of peptide in 3 groups of 6 to 8-week-old female B6 mice; Group 1 was immunized using our standard EAE protocol in B6 mice; Group 2 was immunized using an EAE protocol for HLA-DR2b transgenic mice; Group 3 was immunized using a myelin peptide cocktail. **(C)** EAE immunization protocols used for immunization of humanized B2m-NOG mice. In Experiment 1, PBMC B2m-NOG mice were immunized with the myelin peptide cocktail at 200 μg/peptide, followed by a repeat boost immunization 7 days later. In Experiment 2, DR15 MS2-5 mice were immunized once with the myelin peptide cocktail at 100 μg/peptide.

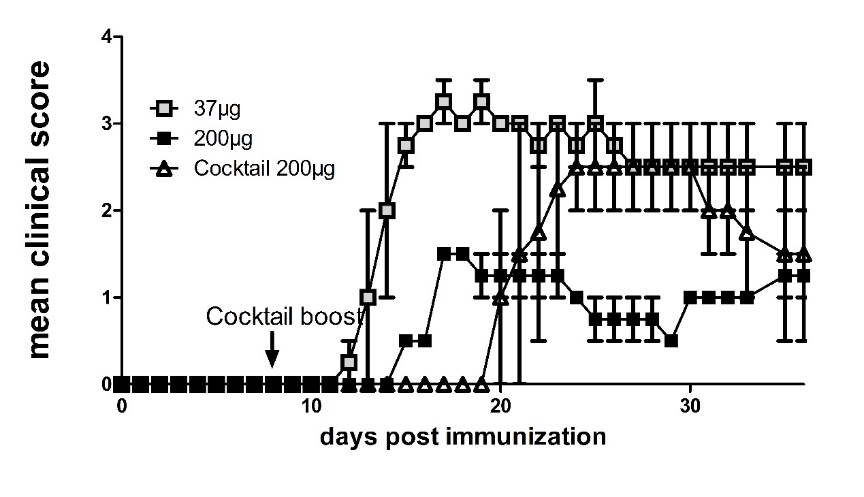
**A: Clinical scores in EAE tests in B6 mice**

**B: EAE peptides in immunization tests in B6 mice**

| **Myelin peptides** | **Group 1**  **B6 (n=2)** | **Group 2**  **B6 (n=2)** | **Group 3**  **B6 (n=2)** |
| --- | --- | --- | --- |
| mMOG35-55 | 37μg | 200μg x 2 | 200μg x 2 |
| hMOG35-55 | - | - | 200μg x 2 |
| MBP83-99 | - | - | 200μg x 2 |
| MOG1-20 | - | - | 200μg x 2 |

**C: Myelin peptides for immunization of PBMC B2m-NOG mice**

| **Myelin peptides** | **Experiment 1**  **PBMC B2m-NOG**  **(DR13 MS, DR15 MS1, DR15 HI)** | **Experiment 2**  **PBMC B2m-NOG**  **(DR15 MS2-5)** |
| --- | --- | --- |
| mMOG35-55 | 200μg x 2 | 100μg x 1 |
| hMOG35-55 | 200μg x 2 | 100μg x 1 |
| MBP83-99 | 200μg x 2 | 100μg x 1 |
| MOG1-20 | 200μg x 2 | 100μg x 1 |
