## Supplementary Table 2 for "Spontaneous human CD8 T cell and EAE-inducible human CD4/CD8 T cell lesions in the brain and spinal cord of HLA-DRB1*15-positive multiple sclerosis PBMC humanized mice"

**Supplementary table 2: Analyses of peripheral blood samples from MS and healthy PBMC donors. (A)** Panel of human immune cell markers used for flow cytometry. **(B)** Immunoprofiling of fresh peripheral blood from donors by flow cytometry using antibody panel in A. (**C)** Clinical interpretation of ELISA results for detection of EBV antibodies in plasma from donors. *****Anomalous result of finding positive levels in both IgG and IgM anti-EBNA1 antibody classes can be a sign of an autoimmune disease.

**A: Panel of human immune cell markers used for flow cytometry**

| **Immune cell marker-fluorochrome** | **Major cell populations identified** |
| --- | --- |
| **hCD45** BV-510 (clone 2D1) | Pan-leukocytes |
| **hCD3** APC/Cy7 (clone HIT3a) | Total T lymphocytes |
| **hCD4** PE/Cy7 (H7) (clone A161A1) | T helper lymphocytes |
| **hCD8** PerCP (clone SK1) | T cytotoxic lymphocytes |
| **hCD19** PE (clone 4G7) | B lymphocytes |
| **hCD56** FITC (clone 5.1H11) | Natural killer cells |
| **hCD66b** APC* (clone G10F5) | Neutrophils |
| **hCD14** BV421 (clone HCD14) | Monocytes, macrophages, dendritic cells |

***** For screening mouse blood this marker was replaced by **mCD45** APC (clone 30-F11)

**B: Immunoprofiling of fresh donor blood by flow cytometry**

|  | **% nucleated cells** | **% hCD45 cells** | | | | | | |
| --- | --- | --- | --- | --- | --- | --- | --- | --- |
| **Donor** | **hCD45** | **hCD3** | **hCD4** | **hCD8** | **hCD19** | **hCD56** | **hCD66b** | **hCD14** |
| **DR13 MS** | 22.6 | 41.4 | 23.4 | 13.8 | 10.7 | 5.4 | 36.6 | 2.2 |
| **DR15 MS1** | 80.6 | 17.1 | 12.4 | 3.8 | 5.0 | 4.6 | 64.0 | 6.6 |
| **DR15 HI** | 84.9 | 34.1 | 22.1 | 9.5 | 3.3 | 6.2 | 48.1 | 3.8 |
| **DR15 MS2** | 97.6 | 30.7 | 17.9 | 10.6 | 6.6 | 3.9 | 50.5 | 7.8 |
| **DR15 MS3** | 98.8 | 28.1 | 18.3 | 9.9 | 7.5 | 3.5 | 50.7 | 7.7 |
| **DR15 MS4** | 99.4 | 23.6 | 15.6 | 8 | 6.1 | 2.3 | 60.1 | 5.3 |
| **DR15 MS5** | 99.6 | 16.8 | 12 | 3.8 | 6.3 | 2.5 | 65.8 | 5.7 |

**C: Clinical interpretation of EBV antibody results**

| **VCA EBV IgG** | **VCA EBV IgM** | **VCA EBV IgA** | **EA(D) EBV IgG** | **EBNA1 EBV IgG*** | **EBNA1 EBV IgM*** | **EBV infection stages** |
| --- | --- | --- | --- | --- | --- | --- |
| - | - | - | - | - | - | Seronegative |
| - | + | + | - | - | + | Primoinfection (early) |
| +  low avidity | + | + | + or - | - | + | Primoinfection |
|  | + | - | + or - | - | + |  |
|  | - | + | + or - | - | - |  |
| +  high avidity | + | - | + or - | + | - | Suspect recent or active reactivation |
|  | - | + | + or - | + | - |  |
|  | - | - | ++ | + | - |  |
| +  high avidity | - | - | - | + | - | Seropositive without symptoms of active infection |

*Individuals positive for both EBNA1 EBV IgG and EBNA1 EBV IgM antibodies are characterized as anomalous EBV reactivation.
