## SupplementaryTable 3 for "Spontaneous human CD8 T cell and EAE-inducible human CD4/CD8 T cell lesions in the brain and spinal cord of HLA-DRB1*15-positive multiple sclerosis PBMC humanized mice"

**Supplementary table 3: Optimization of human PBMC isolation protocol.** Three different PBMC isolation protocols were compared for the preparation of cells for transplantation into B2m-NOG mice. Specifically, PBMC were isolated from peripheral blood samples from a healthy individual, fresh on the day of transplantation (protocol 1), after dilution of fresh blood with culture medium and 3 days’ storage at 4°C (protocol 2), freshly isolated, frozen, thawed and cultured for 24 h (protocol 3).

| **PBMC preparation protocol** | **PBMC yield (10ml blood)** | **% nucleated cells** | **% hCD45 cells** | | | | | | |
| --- | --- | --- | --- | --- | --- | --- | --- | --- | --- |
|  |  | **hCD45** | **hCD3** | **hCD4** | **hCD8** | **hCD19** | **hCD56** | **hCD66b** | **hCD14** |
| 1) Blood, PBMC prepared fresh | 10-13 x 10^6^ | 91.8 | 17.0 | 11.0 | 4.7 | 2.8 | 0.5 | 68.2 | 5.6 |
| 2) Blood, diluted 1:1 Glutamax, stored 72h at 4oC | 7-9 x 10^6^ | 92 | 41.2 | 17.2 | 24.0 | 7.7 | 4.8 | 30.9 | 3.3 |
| 3) PBMC, frozen/thawed, cultured for 24h | 0.5 x 10^6^ | 93.4 | 36.5 | 14.0 | 19.6 | 10.9 | 0.6 | 25.7 | 0.8 |
