## Supplementary Table 4 for "Spontaneous human CD8 T cell and EAE-inducible human CD4/CD8 T cell lesions in the brain and spinal cord of HLA-DRB1*15-positive multiple sclerosis PBMC humanized mice"

**Supplementary table 4: Human immune cell engraftment in PBMC humanized B2m-NOG mouse splenocyte analysis:** Analysis of splenocytes recovered from non-immunized and EAE-immunized (+) PBMC B2m-NOG mice at sacrifice on day post transplantation 42 (dpt 42) by flow cytometry, showing engraftment by human CD45^+^ leukocyte populations.

| **PBMC donor** | **EAE immunization** | **% hCD45 cells** | | |
| --- | --- | --- | --- | --- |
|  |  | **% hCD4^+^**  **T cells** | **% hCD8^+^**  **T cells** | **% hCD19^+^ B cells** |
| **Experiment 1** | | | | |
| **DR13 MS** | - | 50.53 ± 2.77 | 15.79 ± 3.86 | 28.47 ± 2.68 |
|  | + | 45.02 ± 3.82 | 13.49 ± 2.16 | 35.14 ± 3.73 |
| **DR15 MS1** | - | 45.73 ± 2.14 | 44.93 ± 2.53 | 1.78 ± 0.71 |
|  | + | 55.48 ± 3.99 | 39.3 ± 3.91 | 1.65 ± 0.30 |
| **DR15 HI** | - | 64.37 ± 5.67 | 24.4 ± 6.95 | 2.61 ± 0.16 |
|  | + | 72.64 ± 3.04 | 20.44 ± 1.79 | 3.23 ± 1.56 |
| **Experiment 2** | | | | |
| **DR15 MS2** | - | 59.63 ± 12.19 | 32.44 ± 11.96 | 2.80 ± 2.22 |
|  | + | 77.2 ± 6.09 | 15.46 ± 5.99 | 4.15 ± 1.92 |
| **DR15 MS3** | - | 63.2 ± 1.76 | 17.83 ± 5.69 | 15.22 ± 6.43 |
|  | + | 83.38 ± 2.79 | 9.86 ± 2.26 | 4.00 ± 0.87 |
| **DR15 MS4** | - | 68.9 ± 6.30 | 23.13 ± 4.71 | 1.25 ± 0.91 |
|  | + | 77.83 ± 5.97 | 16.69 ± 4.30 | 1.36 ± 1.04 |
| **DR15 MS5** | - | 71.00 ± 3.72 | 20.93 ± 4.98 | 6.71 ± 1.00 |
|  | + | 70.43 ± 2.79 | 22.53 ± 4.10 | 5.13 ± 1.43 |
