## Supplementary Table 5 for "Spontaneous human CD8 T cell and EAE-inducible human CD4/CD8 T cell lesions in the brain and spinal cord of HLA-DRB1*15-positive multiple sclerosis PBMC humanized mice"

**Supplementary table 5:** Comparative flow cytometry analysis of (m) CD11b^+^ myeloid cell subpopulations in the peripheral blood of non-PBMC-engrafted mouse strains. Analysis of blood mCD11b^+^ myeloid cell subpopulations in non-immunized (naïve) B2m-NOG (n=4), NOD-*scid* (n=2) and C57BL/6 (B6) (n=5), as well as in groups of EAE immunized NOD-*scid* (n=2) and B6 (n=5) mice at dpi 8. EAE immunization was performed by s.c. tail-base injection of CFA emulsion supplemented with H37Ra and without myelin peptides, followed by 2 injections of *Bordetella pertussis* toxin, as described in the Material and Methods.

| **Mouse strain**  **(non-PBMC-engrafted)** | **% mCD11b^+^** | **%Ly6C^hi^ of mCD11b^+^** | **%Ly6G^+^ of mCD11b^+^** |
| --- | --- | --- | --- |
| **B2m-NOG** | | | |
| Naïve | 96.38 ± 0.90 | 7.20 ± 1.98 | 74.13 ± 7.11 |
| **NOD-*scid*** | | | |
| Naïve | 87,6 | 12,7 | 54,7 |
| CFA immunization (dpi 8) | 91.8 ± 3.61 | 13.7± 0.42 | 62.7 ± 4.31 |
| **C57BL/6** | | | |
| Naïve | 10.46 ± 0.38 | 13.15 ± 0.53 | 36.45 ± 1.10 |
| CFA immunization (dpi 8) | 40.58 ± 2.76 | 15.02 ± 0.71 | 66.24 ± 1.22 |
